# Fructose-2,6-bisphosphate restores TDP-43 pathology-driven genome repair deficiency in motor neuron diseases

**DOI:** 10.1101/2024.11.13.623464

**Authors:** Anirban Chakraborty, Joy Mitra, Vikas H. Maloji Rao, Manohar Kodavati, Santi M. Mandal, Satkarjeet K. Gill, Sravan Gopalkrishnashetty Sreenivasmurthy, Velmarini Vasquez, Mikita Mankevich, Ludo Van Den Bosch, Ralph M. Garruto, Ian Robey, Balaji Krishnan, Gourisankar Ghosh, Muralidhar L. Hegde, Tapas Hazra

## Abstract

TAR DNA-binding protein 43 (TDP-43) proteinopathy plays a critical role in neurodegenerative diseases, including amyotrophic lateral sclerosis (ALS) and frontotemporal dementia (FTD). We recently reported that TDP-43 plays an essential role in DNA double-strand break (DSB) repair via the non-homologous end-joining (NHEJ) pathway. Here, we provide evidence that the brain of patients with ALS exhibit persistent DNA damage in the transcribed regions of the genome. While investigating the mechanistic basis, we found that the activity of polynucleotide kinase 3’-phosphatase (PNKP) was severely impaired in the nuclear extracts of patient brains and TDP-43-depleted cells. PNKP is a key player in DSB repair within the transcribed genome, where its 3’-phosphate termini processing activity is crucial for preventing persistent DNA strand breaks and neuronal death. The inactivation of PNKP was due to the reduced level of its interacting partner, phosphofructo-2-kinase fructose 2,6 bisphosphatase (PFKFB3), and its biosynthetic product, fructose-2,6-bisphosphate (F2,6BP), an allosteric modulator of glycolysis. Recently, we have demonstrated that F2,6BP acts as a positive modulator of PNKP activity *in vivo*. Furthermore, F2,6BP supplementation in cultured ALS patient-derived neural progenitor stem cells (NPSCs) reduced the toxic aggregation of polyubiquitinated TDP-43 and cytosolic pTDP-43 (S409/410). Notably, F2,6BP supplementation restored the PNKP activity in the nuclear extracts from autopsied ALS/FTD brain tissues and patient iPSC-derived NPSCs harboring TDP-43 mutations. Importantly, F2,6BP administration significantly restored the genome integrity and motor phenotypes in a Drosophila model of ALS-TDP-43. Collectively, these findings underscore the therapeutic potential of F2,6BP in TDP-43 pathology-associated motor neuron diseases.

## Introduction

Amyotrophic lateral sclerosis (ALS) and frontotemporal dementia (FTD) are neurodegenerative diseases predominantly marked by dysfunction and toxicity of TAR DNA-binding protein of 43 kDa (TDP-43), which drives the progressive degeneration of motor neurons (1–5). TDP-43 pathology in these diseases is characterized by the loss of nuclear TDP-43 and its accumulation in the cytoplasm, disrupting cellular pathways in motor neurons (6, 7). Our recent studies demonstrated that TDP-43 plays a key role in the classical non-homologous end-joining (C-NHEJ) pathway, which is primarily responsible for repairing DNA double-strand breaks (DSBs) in post-mitotic neurons. Upon DNA damage, TDP-43 is rapidly recruited to DSB sites, where it stably interacts with DNA damage response (DDR) and NHEJ factors, facilitating the recruitment of classical NHEJ factors, including the break-sealing XRCC4-DNA Ligase 4 (Lig4) complex in induced pluripotent stem cell (iPSC)-derived motor neurons (5, 8). We also showed that mislocalization of ALS-linked mutant TDP-43 significantly impaired NHEJ-mediated DSB repair, leading to the accumulation of DNA damage, cellular senescence, and increased neuroinflammation (9). Notably, the compromised NHEJ repair in the TDP-43 proteinopathy is associated with reduced recruitment of the XRCC4-Lig4 complex at DSB sites (5, 8). However, the precise mechanism by which TDP-43 regulates C-NHEJ remains unclear.

In ALS and FTD, enhanced production of reactive oxygen species (ROS) and persistent neuroinflammation can lead to oxidative genome damage and DNA single-strand breaks (SSBs) and double-strand breaks (DSBs) (10–14). Polynucleotide kinase 3’-phosphatase (PNKP) is a bifunctional DNA end-processing enzyme with 3’-phosphatase and 5’-kinase activities (15, 16), and a major 3’-phosphatase in mammalian cells that converts repair-incompetent broken DNA ends to ligatable ends, a crucial step in DSB repair. It is involved in several DNA repair pathways, including base excision repair (BER), SSB, and C-NHEJ-mediated DSB repair in mammalian cells (17–22). Importantly, functional loss of PNKP in neuronal cells has been linked to several neuropathological conditions (23–29). Additionally, we have demonstrated that PNKP plays an essential role in the preferential repair of actively transcribed genomes via transcription-coupled SSB (TC-SSBR) and NHEJ (TC-NHEJ) repair pathways (17, 21, 30–35). Notably, our studies reveal that the 3′-phosphatase activity of PNKP is severely compromised in the two most prevalent polyQ neurodegenerative diseases - Huntington’s disease (HD) (30, 36) and spinocerebellar ataxia type 3 (SCA3) (31, 36).

Several studies provided strong mechanistic evidence linking metabolic reprogramming and DNA repair activity through the direct involvement of metabolic enzymes and metabolites (37–39). We recently showed that 6-phosphofructo-2-kinase fructose-2,6-bisphosphatase 3 (PFKFB3), the only member in the PFKFB family (PFKFB1-4) present in the nucleus (40, 41), associates with and regulates PNKP functionality in mammalian cells. PFKFB3 converts fructose-6-phosphate (F6P) to fructose-2,6-bisphosphate (F2,6BP), a key inducer of glycolysis. Notably, the levels of both PFKFB3 and its product, F2,6BP, are significantly lower in the nuclear extract (NE) of autopsied HD and SCA3 patients’ brain tissues. Interestingly, supplementation of F2,6BP in HD mouse-derived striatal neurons and HD flies restores nuclear and mitochondrial genome integrity and functionality (36, 42), suggesting F2,6BP to be a positive regulator of PNKP activity *in vivo*. Reduced levels of PFKFB3 are therefore linked to the decreased PNKP activity in HD and SCA3, connecting metabolic processes to DNA repair (31, 36).

TDP-43 plays a crucial role in C-NHEJ, interacting with other NHEJ factors, such as XRCC4-Lig4, which also associate with PNKP and PFKFB3 (5, 17). Therefore, we reasoned that these proteins might be part of the same molecular complex involved in DSB repair. PNKP interactome studies showed that TDP-43 is associated with PNKP, PFKFB3, Lig4 and other repair proteins, forming a complex involved in TC-NHEJ. These observations led us to investigate the role of PNKP and PFKFB3 in ALS and FTD models showing TDP-43 pathology. We observed a near-complete loss of 3’-phosphatase activity of PNKP, but no change in the protein level, under TDP-43-associated neurodegenerative conditions, correlating with TDP-43 mislocalization and persistent DNA damage. Furthermore, we observed reduced levels of PFKFB3 and F2,6BP in ALS/FTD patient tissues, indicating potential disruptions in the glycolytic pathway that may exacerbate genomic instability associated with TDP-43 pathology. Importantly, such PNKP-mediated repair deficiency could be restored by exogenous supplementation of F2,6BP in extracts isolated from ALS/FTD patients’ brain tissues and patient-derived NPSCs. Supplementation of F2,6BP reduced pathogenic phosphorylation and ubiquitination of TDP-43, resulting in a significant reduction of aggregates of TDP-43 in ALS patient-derived NPSCs. The fruit fly, *Drosophila melanogaster*, has emerged as a powerful *in vivo* system to dissect the complex cellular and molecular mechanisms underlying neurodegenerative diseases, owing to its sophisticated genetic toolkit and the high conservation of fundamental biological pathways with humans (43). Notably, F2,6BP supplementation also significantly rescued motor deficiency in a *Drosophila* model of ALS harboring TDP-43^Q331K^ mutation, a well-established model that recapitulates key features of the human disease, including progressive, age-dependent motor decline and neurodegeneration (44, 45). Overall, our findings highlight the therapeutic potential of F2,6BP in ALS, FTD and other neurodegenerative diseases linked to TDP-43 pathology via restoring DNA repair.

## Methods

### Cell culture and stable cell line generation

Human embryonic kidney HEK293 (ATCC) cells were cultured in Dulbecco’s modified Eagle’s medium (DMEM) with high glucose, supplemented with 10% fetal bovine serum (FBS) and 100 U/ml penicillin-streptomycin in a humidified chamber at 37°C and 5% CO2. To generate the PNKP-FLAG-expressing stable HEK293 cell line, the coding DNA sequence of human PNKP (GenBank BC033822.1) was re-cloned to pFLAG-cDNA (Invitrogen/Life Technologies) between the CMV promoter and FLAG-tag encoding sequences at *Hin*dIII and *Bam*HI sites. The gene in the newly constructed plasmid carried the natural Kozak sequence and nucleotides 97–1,669 of PNKP cDNA. The region containing Kozak-PNKP-FLAG was then transferred to pcDNA3.1-Hygro (Invitrogen/Life Technologies) within the *Hin*dIII-*Xb*aI unique vector sites to generate the final plasmid, which was then used for generating a stable cell line resistant to hygromycin (Hygro), following standard procedures (17). Ectopic expression of PNKP was confirmed by western blotting of whole-cell extract using anti-FLAG Ab (Sigma, F1804).

### siRNA transfection

TDP-43 depletion was carried out in HEK293 cells using siRNAs (80 nM; transfected twice on consecutive days) against the *TARDBP* gene. The cells were transfected with control siRNA (siControl) or TDP-43-specific siRNA (siTDP-43; Thermo Fisher; Silencer Select; 4392420) using Lipofectamine 3000 (Invitrogen) for 6 h in optiMEM (Gibco; reduced serum media, 11058021), followed by replacing the medium with DMEM-High Glucose complete medium. Nuclear extracts (NE) were prepared from the harvested cells (72-96 h post-transfection) to examine the depletion of individual proteins by immunoblot (IB) analysis using specific antibodies.

### Neural Progenitor Stem Cell (NPSC) induction and motor neuron differentiation

Control and TDP-43 mutant induced pluripotent stem cells (iPSCs) were cultured on basement membrane matrix Geltrex LDEV-Free-coated Petri dishes using 1X Essential 8 media (Gibco, A1517001) and maintained in a humidified incubator at 37°C in 5% CO_2_. To derive NPSCs, PSC neural induction medium (Gibco A1647801) was used, as per the manufacturer’s protocol. The process involved replacing the Essential 8 media with PSC neural induction media approximately 24 h after sub-plating those iPSCs. This media was maintained for 7 days. The first passage (P0) NPSCs were then transferred onto Geltrex (Gibco, A1413201) coated 6-well plates and cultured in StemPRO neural stem cell SFM media (Gibco, A1050901). Neural induction efficiency was assessed at the third passage by immunofluorescence (IF) staining.

ALS-linked TDP-43^G287S^ mutant and its CRISPR/Cas9-corrected isogenic TDP-43^G287G^ iPSCs, as well a FTD-related TDP-43^A382T^ iPSCs also containing the C9ORF72 repeats were kind gifts from our collaborators Drs. Van Damme and Van Den Bosch (University of Leuven, Belgium). Details about these iPSC lines and the protocols used to culture these cells were published before (46). Control and ALS patient-derived TDP-43^G298S^ mutant iPSC (NINDS, NH50216) lines were purchased from NIH-NINDS (47). Engineered homozygous TDP-43^Q331K^ mutant (Jax, JIPSC1064) and its isogenic control (Jax, JIPSC1104) lines were purchased from Jax iPSC.

Motor neurons were generated from engineered iPSCs with TDP-43^Q331K^ mutation and its isogenic control line. The differentiation process followed established protocols with some modifications (48, 49). In brief, iPSC clones were transferred from a 60-cm dish to a T-25 flask filled with neuronal basic medium. The medium consisted of a mixture of 50% Neurobasal medium (Gibco, 21103049) and 50% DMEM/F12 medium (Gibco, 21331020), supplemented with N2 and B27 supplements without vitamin A. Collagenase type IV digestion was performed to facilitate suspension of the iPSC clones. Afterward, the suspended cell spheres were subjected to a series of incubations. Initially, they were treated with various inhibitors, including 5 μM ROCK inhibitor (Y-27632), 40 μM TGF-β inhibitor (SB 431524), 0.2 μM bone morphogenetic protein inhibitor (LDN-193189), and 3 μM GSK-3 inhibitor (CHIR99021). This was followed by incubation in a neuronal basic medium containing 0.1 μM retinoic acid (RA) and 500 nM Smoothened Agonist (SAG) for 4 days. Subsequently, the cell spheres were incubated for 2 days in a neuronal basic medium containing RA, SAG, 10 ng/ml brain-derived neurotrophic factor (BDNF), and 10 ng/ml glial cell-derived neurotrophic factor (GDNF). To dissociate the cell spheres into single cells, they were exposed to a neuronal basic medium containing trypsin (0.025%)/DNase in a water bath at 37°C for 20 min. Afterward, the cells were pipetted into a medium containing trypsin inhibitor (1.2 mg/ml) to maintain the viability. Following cell counting, a specific number of cells were seeded onto dishes or chamber slides coated with 20 μg/ml Laminin. These cells were incubated for 5 days in a basic neuronal medium containing RA, SAG, BDNF, GDNF, and 10 μM Inhibitor of γ-secretase (DAPT). Subsequently, the medium was switched to one containing BDNF, GDNF, and 20 μM DAPT for an additional 2 days. For motor neuron maturation, the cells were cultured in a medium containing BDNF, GDNF, and 10 ng/ml ciliary neurotrophic factor (CNTF) for a period exceeding 7 days.

### ALS, FTD and Guamanian ALS patient brain samples

The cortical brain samples of sporadic ALS, FTD, Guamanian ALS and their non-neurological controls were obtained respectively from the Veterans Affairs Brain Biorepository (VABB) and Binghamton University Brain Biorepository (see the cohort lists in *Supplementary* **Table S1**). For preparing the nuclear and cytosolic fractions of protein samples, ∼30 mg of brain tissue powder was used for each sample, followed by stepwise fractionations of protein extracts using methods described in later sections. For PNKP activity assays, nuclear extracts were freshly prepared and subjected to the assay immediately for reliable and reproducible outcomes. For the extraction of genomic DNA from these tissue samples, about 20 mg of tissue powder was used for each sample, and the genomic DNA was extracted using commercial kits, as described later.

### ALS mouse brain tissue

In this study, we have utilized an established mouse model of ALS carrying an endogenous murine Tdp-43^ΔNLS^ mutation. The Tdp-43^ΔNLS^ variant was expressed under a motor neuron-specific gene promoter, Mnx1 (Hb9), allowing the Tdp-43 pathology to initiate as neurons differentiate with aging and exhibited ALS-like pathological phenotypes at one year of age (9). At this time point, mice were humanely sacrificed, and their brain cortices were harvested for biochemical assays.

### Immunoblotting (IB)

For IB analysis, cell lysates were prepared with 1x RIPA buffer (Millipore, 20-188) containing the protease inhibitor cocktail (Roche, 11836170001). Protein concentration was measured using 1x Bradford reagent (Biorad) in NanoDrop instrument.

For IB with insoluble fractionates, the pellet from the soluble fractionation of each sample was dissolved in an equal volume of SDS buffer containing 2% SDS, 50 mM Tris-HCl pH 8.0, and 10% glycerol, followed by sonication (50). Probing with GAPDH in soluble fractions was used to confirm the uniformity in the amounts of starting cell pellets.

Then, 20 µg protein or an equal volume of sample was loaded into NuPAGE 4–12% Bis-Tris precast gels (Invitrogen) for electrophoretic resolution. After transferring to nitrocellulose membranes, the separated proteins were incubated with target primary and respective secondary antibodies, and the protein signals were detected by adding chemiluminescence reagents (LI-COR) and visualized by LI-COR Odyssey imaging system.

### Co-immunoprecipitation (Co-IP)

Approximately 30 mg of brain tissues from ALS/FTD post-mortem patients or healthy controls (*Supplementary* **Table S1**) were homogenized with 4 volumes of ice-cold homogenization buffer [0.25 M sucrose, 15 mM Tris-HCl pH 7.9, 60 mM KCl, 15 mM NaCl, 5 mM EDTA, 1 mM EGTA, 0.15 mM spermine, 0.5 mM spermidine, 1 mM dithiothreitol (DTT), 0.1 mM phenylmethylsulfonyl fluoride (PMSF), and protease inhibitors (EDTA-free; Roche)]. Following homogenization, the homogenate was incubated on ice for 15 min and centrifuged at 1,000×g to obtain the cell pellet. Cells were lysed in buffer A [10 mM Tris-HCl pH 7.9, 0.34 M sucrose, 3 mM CaCl_2_, 2 mM magnesium acetate, 0.1 mM EDTA, 1 mM DTT, 0.5% Nonidet P-40 (NP-40) and 1x protease inhibitor cocktail (Roche)] and centrifuged at 3,500×g for 15 min. Nuclear pellets were washed with buffer A without NP-40 and then lysed in buffer B [20 mM HEPES pH 7.9, 3 mM EDTA, 10% glycerol, 150 mM potassium acetate, 1.5 mM MgCl_2_, 1 mM DTT, 0.1% NP-40, 1 mM sodium orthovanadate (vanadate) and 1x protease inhibitors] by homogenization. Supernatants were collected after centrifugation at 15,000×g for 30 min and DNA/RNA in the suspension was digested with 0.15 U/μl benzonase (Novagen, 70-746-3) at 37°C for 1 h. The samples were centrifuged at 20,000×g for 30 min, and the supernatants collected as NEs. Co-IP was performed using anti-PNKP (BioBharati Life Science, BB-AB0105) antibody with Protein A/G PLUS agarose beads (Santa Cruz Biotechnology, sc2003) overnight, followed by four washes with Wash buffer (20 mM HEPES pH 7.9, 150 mM KCl, 0.5 mM EDTA, 10% glycerol, 0.25% Triton-X-100 and 1x protease inhibitors) and eluted with Laemmli Sample Buffer (Bio-Rad; final concentration 1x). The immunoprecipitates were tested for the interacting proteins using appropriate Abs [PFKFB3 (GeneTex, GTX108335), Lig 4 (GeneTex, GTX108820), and TDP-43 (Protein tech, 10782-2-AP)].

### Assay of 3D-phosphatase activity of PNKP

The 3’-phosphatase activity of PNKP in 1 µg of NE of post-mortem ALS/FTD and age-matched patients’ cortical tissues or with purified recombinant PNKP (2 ng) was assessed, as described previously (36). Five pmol of the radiolabeled substrate was incubated at 37°C for 15 min in buffer A (25 mM Tris-HCl pH 8.0, 100 mM NaCl, 5 mM MgCl_2_, 1 mM DTT, 10% glycerol and 0.1 μg/μl acetylated BSA). Five pmol of non-radiolabeled substrate was used as a cold substrate. For *in vitro* PNKP restoration, similar assays were done after incubation of F2,6BP or F6P/F1,6BP (as controls) with the NE for 15 min. The radioactive bands were visualized in PhosphorImager (GE Healthcare) and quantified using ImageQuant software.

### Enzymatic preparation of F2,6BP

Enzymatic preparations of F2,6BP were conducted, as we described earlier (36).

### Estimation of F2,6BP in the patient brain NE

To quantify F2,6BP, the F6P assay kit (Sigma-Aldrich) was used to measure the amount of F6P before and after treating the crude NE with 0.1M HCl and incubating at 37°C for 30 min. Acid treatment converts F2,6BP into F6P. The assay was performed using the manufacturer’s protocol, which uses NADH-linked luciferase bioluminescence (36).

### Long Amplicon Quantitative PCR (LA-qPCR)

Genomic DNA was extracted from cultured cells, autopsied brain tissues or adult *Drosophila* using the Genomic tip 20/G kit (Qiagen) per manufacturer’s protocol, to ensure minimal DNA oxidation during the isolation steps. The DNA was quantified by Pico Green (Molecular Probes) in a black-bottomed 96-well plate and gene-specific LA-qPCR assays were performed as described earlier (36) using Long Amp Taq DNA Polymerase (New England Biolabs). POLB and RNAPII (12.1 and 11.3 kb) were amplified from experimental and control samples using appropriate oligos. Enolase/NeuroD (transcribed genes; 6 kb) and MyH2/MyH4 (non-transcribed genes; 6 kb) were amplified from human post-mortem samples. Genomic DNA isolated from adult *Drosophila* (10 flies from each group, male and female mixed) was used for DNA damage analysis. Two genes (CrebB and Neurexin, ∼8 kb) were amplified using appropriate oligos (PMID: 32205441; 39298485). The primers used in this study are detailed in *Supplementary* **Table S2**.

The LA-qPCR reaction was set for all genes from the same stock of diluted genomic DNA (10-15 ng) sample to avoid variations in PCR amplification during sample preparation. The final PCR reaction conditions were optimized at 94°C for 30 s; (94°C for 30 s, 55-60°C for 30 s depending on the oligo annealing temperature, 65°C for 10 min) for 25 cycles; 65°C for 10 min. Since amplification of a small region is independent of DNA damage, a small DNA fragment (∼200-400 bp) from the corresponding gene(s) was also amplified for normalization of amplification of the large fragment. The amplified products were then visualized on gels and quantified using ImageJ software (NIH).

### Immunofluorescence (IF)

For NPSCs and motor neurons, the chamber slides were pre-coated with Geltrex and Matrigel, respectively, to facilitate cell adherence. Fixation of the cells for IF analysis was performed by replacing the media with a mixture of fresh media and 8% paraformaldehyde (PFA) in PBS at a 1:1 ratio, resulting in a final concentration of 4% PFA. Post-fixation, the slides were permeabilized using 0.2% Triton X-100 in 1x PBS, followed by blocking with 5% goat serum-TBS-T (1x TBS with 0.1% Tween-20) to prevent non-specific antibody binding. The cells were then incubated overnight at 4°C with target primary antibodies. Following this step, Alexa Fluor-488 (green) and 647-conjugated secondary antibodies (Thermo Fisher) were incubated for 1 h, and slides were then mounted with coverslips after applying DAPI-containing mounting media (Sigma-Aldrich, USA) to visualize the nuclei. Imaging was performed using a Zeiss Axio Observer 7 microscope or an Olympus Flouview3000 confocal microscope.

The primary antibodies used for various experiments were as follows: rabbit anti-TDP-43 (Proteintech, #10,782–2-AP), mouse anti-TDP-43 (R&D Systems, #MAB77781), rabbit anti-phospho-TDP-43 (S409/410) (Proteintech, #80,007–1-RR), and rabbit anti-ubiquitin (Abcam, #19247).

### Proximity Ligation Assay (PLA)

PLA was conducted to investigate direct protein-protein interactions within cells. Approximately 20,000 cells were seeded per well in 8-well chamber slides for this experiment. After staining, cells underwent a washing step and were then fixed with 4% PFA for 15 min at room temperature. Subsequent steps included permeabilization in 0.2% Triton X-100 in 1x PBS for 10 min at RT, followed by washes in PBS to remove any residual permeabilization agent. The in-situ PLA experiment was performed per the manufacturer’s guidelines, using the DuoLink kit (Sigma-Aldrich, USA). After the PLA procedure, coverslips were mounted using DAPI-containing media (Sigma-Aldrich, USA), and imaging was performed using either a Zeiss Axio Observer 7 microscope or an Olympus Flouview3000 confocal microscope.

### Immunohistochemistry (IHC)

Tissue samples were paraffin-embedded and sliced into 5 μm horizontal sections and mounted on charged glass slides. Slides were dewaxed and autoclaved for 10 min at 121°C in 0.01 M citrate buffer pH 6.0 for antigen retrieval. Immunostaining was performed using overnight incubation at 4°C with mouse anti-TDP-43 (R&D Systems, MAB77781, 1:200), and rabbit anti-phospho-TDP-43 (S409/410) (Proteintech, 80007-1-RR, 1:300). Nuclei were counterstained with DAPI. Slides were imaged in a confocal laser microscope.

### Comet assay

The neutral comet assay was performed using the Comet Assay Kit (Trevigen, 4250-50-K), according to the manufacturer’s protocol, to assess the extent of DNA DSBs in each sample. Briefly, a single-cell suspension was prepared by trypsinization of respective cell types in DPBS buffer, and about 200 cells were smeared in LM Agarose at 37°C in duplicate on each slide. The comet tails were visualized by staining the DNA with SYBR Gold stain under a fluorescence microscope.

### *Drosophila* maintenance and treatment

All *Drosophila* stocks were maintained at 25°C on standard fly food under a 12:12-h light–dark cycle. *Drosophila* strains were purchased from Bloomington Drosophila Stock Center (BDSC, Bloomington, Indiana, USA: UAS-TDP43 Q331K (RRID: BDSC_79590), which expresses human ALS-linked TDP43^Q331K^ mutant. The driver-P{GawB}OK6 (RRID: BDSC_64199) expresses GAL4 primarily in motor neurons. BDSC_79590 flies were crossed to BDSC_64199, and flies eclosed from the crosses were collected in fresh food vials and kept under standard conditions for 1 to 2 d for acclimatization. Flies were separated into two vials for mock buffer and F2,6BP treatment. One BD syringe (1 ml) was filled with mock buffer, and the other was filled with F2,6BP (50 µM), and one drop was administered per day to respective cohorts at the same time of the day for 21 consecutive days.

### Climbing or negative geotaxis assay

The climbing assay was performed as described elsewhere with minor modifications (31, 36). Briefly, experimental flies were anesthetized on ice. A group of 10 flies (male and female mixed) per vial was transferred to a 25 ml sterile glass measuring cylinder. The measuring cylinder was divided into six compartments equally; the lowest compartment was labeled with 1 and the highest compartment was labeled with 6. The measuring cylinder with flies was placed against a white background for better video recording. The cylinder was tapped gently three times to send the flies to the bottom of the cylinder. The climbing time was recorded for 20 s. Five trials were performed for each cohort. The climbing score was calculated at 8 s.

### Statistical Analysis

All data are expressed as mean ± standard deviation (SD). Comparisons among experimental groups were carried out using Student’s t-test, one-way or two-way ANOVA, as appropriate, for significance. Statistical analyses were performed using GraphPad Prism version 10 software. A p-value of less than 0.05 was set as statistically significant.

## Results

### PNKP activity is significantly reduced in ALS and FTD

While our recent studies demonstrated the accumulation of DNA DSBs and a deficiency in NHEJ-mediated DSB repair in ALS brains with TDP-43 proteinopathies (5), the potential link between PNKP and TDP-43 has not yet been investigated in the context of DSB repair. Therefore, we assessed the impact of TDP-43 proteinopathies on the 3’-phosphatase activity of PNKP in ALS and FTD brains [validated by cytosolic mislocalization of total TDP-43 and corresponding levels of pathogenic phosphorylated TDP-43 (pTDP-43, S409/410); *Supplementary* **Figure S1A**] using a duplex oligo-based *in vitro* assay (schematically shown in **Figure 1A)**. We observed a significant inhibition of PNKP activity in the NE isolated from ALS-TDP-43 (Lns 5-7) and FTD-TDP-43 (Lns 8-9) cortical tissues, compared to age-matched healthy controls (Lns 2-4) (**Figure 1B**). We further validated this observation in a second cohort of Guamanian-ALS patients (Lns 6-9) and age-matched healthy control (Lns 2-5) brain cortical tissues and found a significant and comparable reduction (∼4 fold) of PNKP activity, reinforcing the association between TDP-43 pathology and impaired PNKP function (**Figure 1C**). Demographic descriptions of ALS, FTD patients and age-matched non-neurological controls are detailed in *Supplementary* **Table S1**. Furthermore, we quantified relative PNKP levels in the patient brain extracts by immunoblotting (IB) and found them to be comparable to healthy controls (*Supplementary* **Figure S1B**), indicating the loss of PNKP activity to be independent of its protein levels in ALS and FTD brains. These results collectively showed that dysfunction of PNKP is common in ALS, Guam-ALS and FTD cases with TDP-43 proteinopathies.

**Figure 1.**
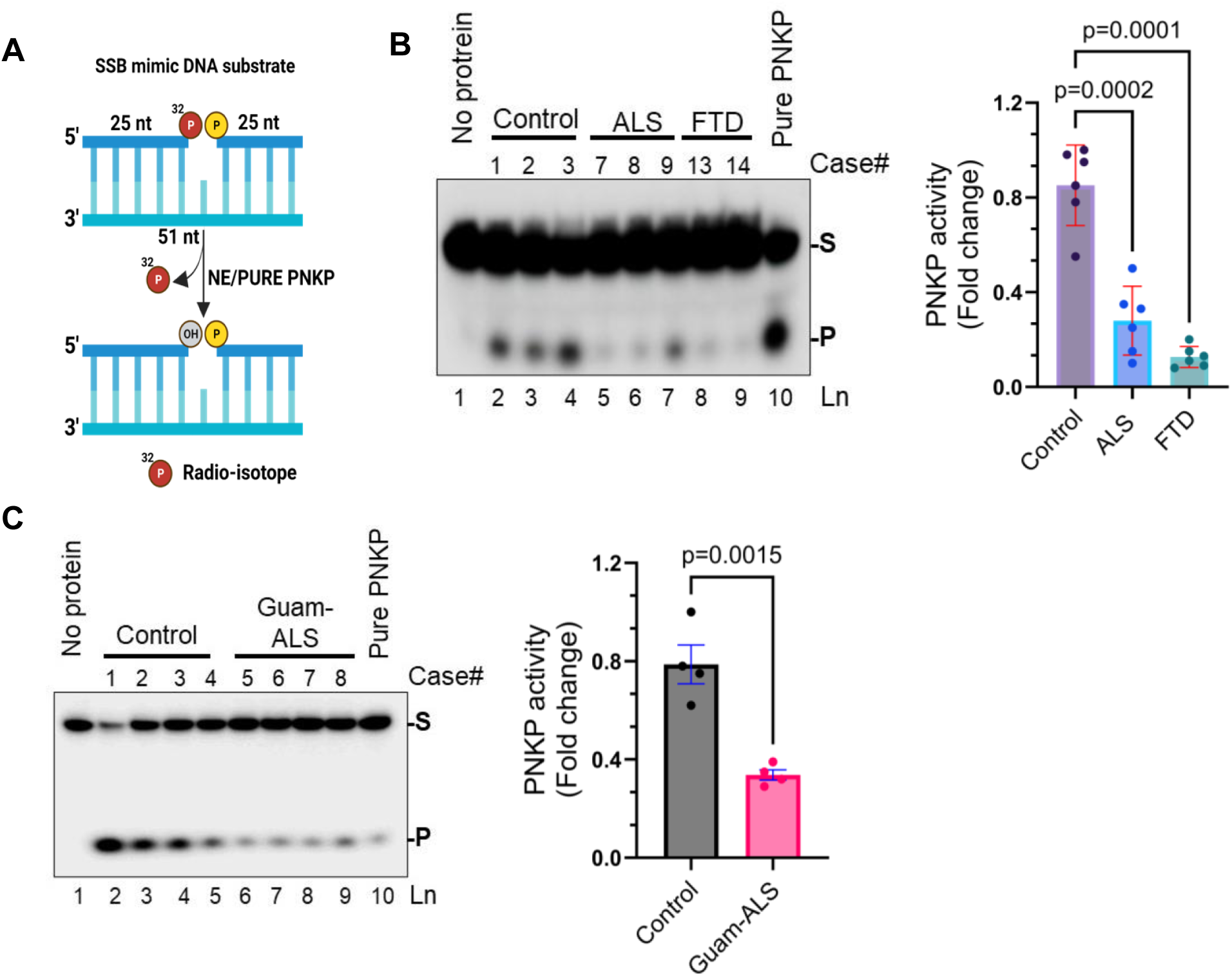
PNKP activity significantly diminishes in sporadic ALS and FTD patient tissues with TDP-43-positive inclusions. **A)** Schematic representation of radio-labeled 32P release by 3’-phosphatase activity of PNKP on a duplex oligonucleotide substrate simulating a DNA single-strand break (SSB). **B)** Representative image demonstrating reduced PNKP activity in the nuclear extracts (NE) of ALS-TDP-43 (Lns 5-7) and FTD-TDP-43 (Lns 8-9) cortical tissues compared to age-matched non-neurological controls (Lns 2-4). Ln 1: substrate only (negative control); Ln 10: purified PNKP (25 fmol) as positive control. The histogram displays quantification of PNKP activities for all samples (N = 6) for the Control, ALS and FTDa **C** groups as fold changes. S: substrate, P: released phosphate. **C)** PNKP activity assay from Guamanian ALS patient brain samples (Lns 6-9) compared to age-matched Guam controls (Lns 2-5). The groups were compared using two-way ANOVA or two-tailed t-tests as appropriate. Error bars represent mean ± SD; significance at p ≤ 0.05; ns = non-significant.

### Downregulation of TDP-43 impairs PNKP activity in cultured cells

To further elucidate the mechanism of how PNKP activity might be regulated by functional loss of TDP-43, we performed siRNA (siTDP-43)-mediated knockdown (KD) of TDP-43 in HEK293 cells, which showed a significant KD of TDP-43 without affecting the PNKP protein level, compared to control siRNA (siControl)-treated cells (**Figure 2A**). Consistently, TDP-43 KD revealed a marked decrease in 3’-phosphatase activity of PNKP in siTDP-43-(Ln 4) compared to siControl-treated (Ln 3) cells (**Figure 2B**). These findings indicate that the loss of TDP-43 directly affects DNA end-processing activity of PNKP, independent of its expression levels, and thereby contributes to the persistent DNA strand breaks in ALS and FTD.

**Figure 2.**
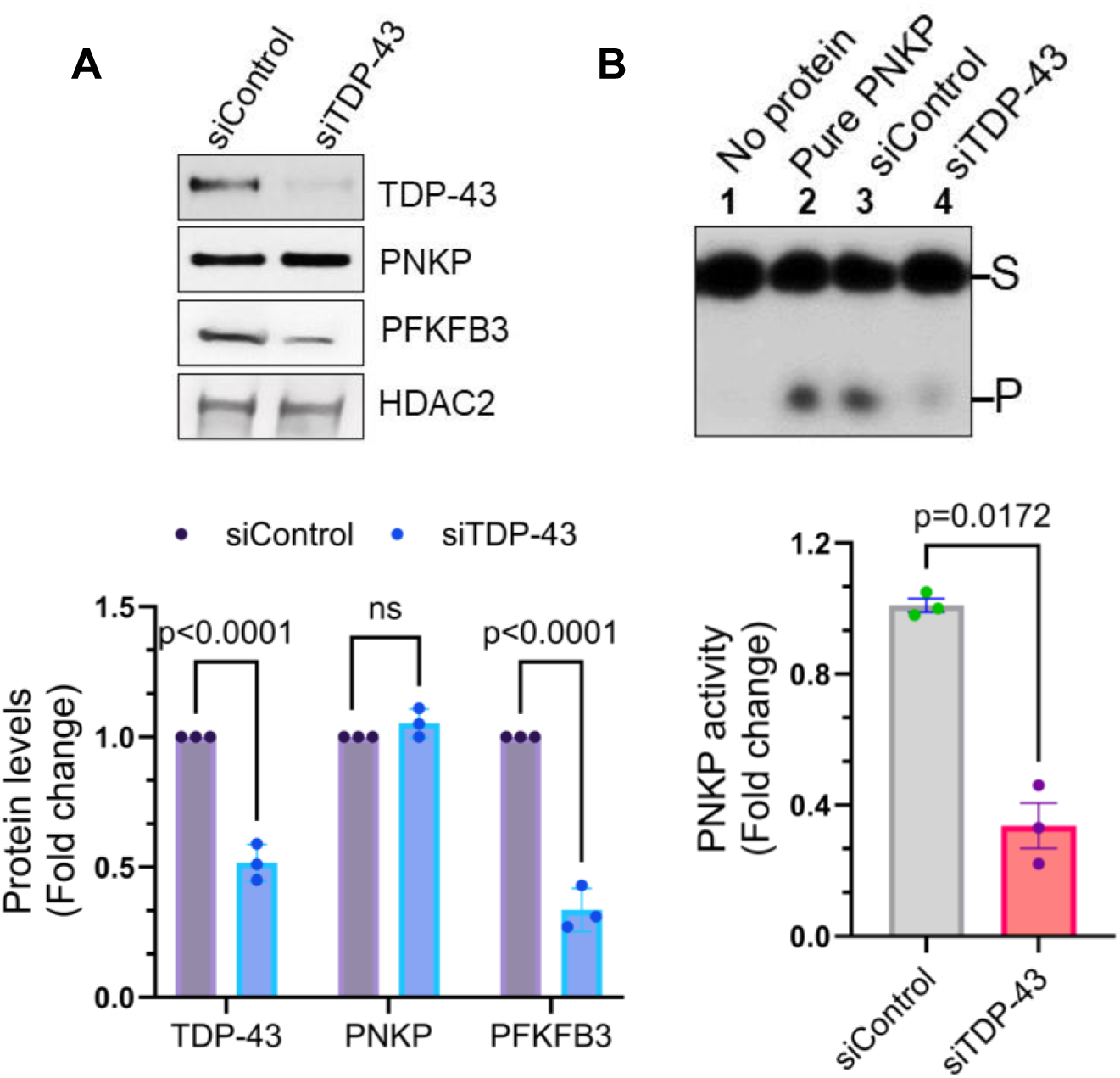
Loss of TDP-43 abolishes PNKP activity in cultured cells. **A)** IB analysis showing levels of TDP-43, PNKP, and PFKFB3 in the NE from HEK293 cells treated with siTDP-43 or siControl. HDAC2 served as the nuclear loading control. The histogram shows quantification of protein levels as fold change, analyzed using two-way ANOVA. **B)** Representative image of 3’-phosphatase activity of PNKP in the NE from HEK293 cells transfected with siControl (Ln 3) or siTDP-43 (Ln 4). Ln 1: substrate alone; Ln 2: purified PNKP (25 fmol) as controls. Quantification of PNKP activity is presented as fold change. Data were analyzed using a Student’s t-test. Error bars represent mean ± SD. The groups were compared using the two-way ANOVA or two-tailed t-tests as appropriate; significance at p ≤ 0.05; ns = non-significant.

### PNKP overexpression rescues TDP-43 KD-associated DNA damage in cells

To determine whether PNKP complementation can mitigate TDP-43 depletion-induced DNA damage, we transfected HEK293 cells, ectopically expressing FLAG-PNKP, with siControl or siTDP-43 and assessed the levels of DSB marker γH2AX across the groups. The results revealed that PNKP overexpression (OE) significantly reduced γH2AX levels in siTDP-43 cells compared to respective control cells (**Figure 3A**). In addition, the neutral comet assay demonstrated a marked reduction in DSB in terms of decreased comet tail moments in PNKP-OE + siTDP-43-treated cells compared to only siTDP-43-transfected cells, indicating improved DNA integrity (**Figure 3B**).

**Figure 3.**
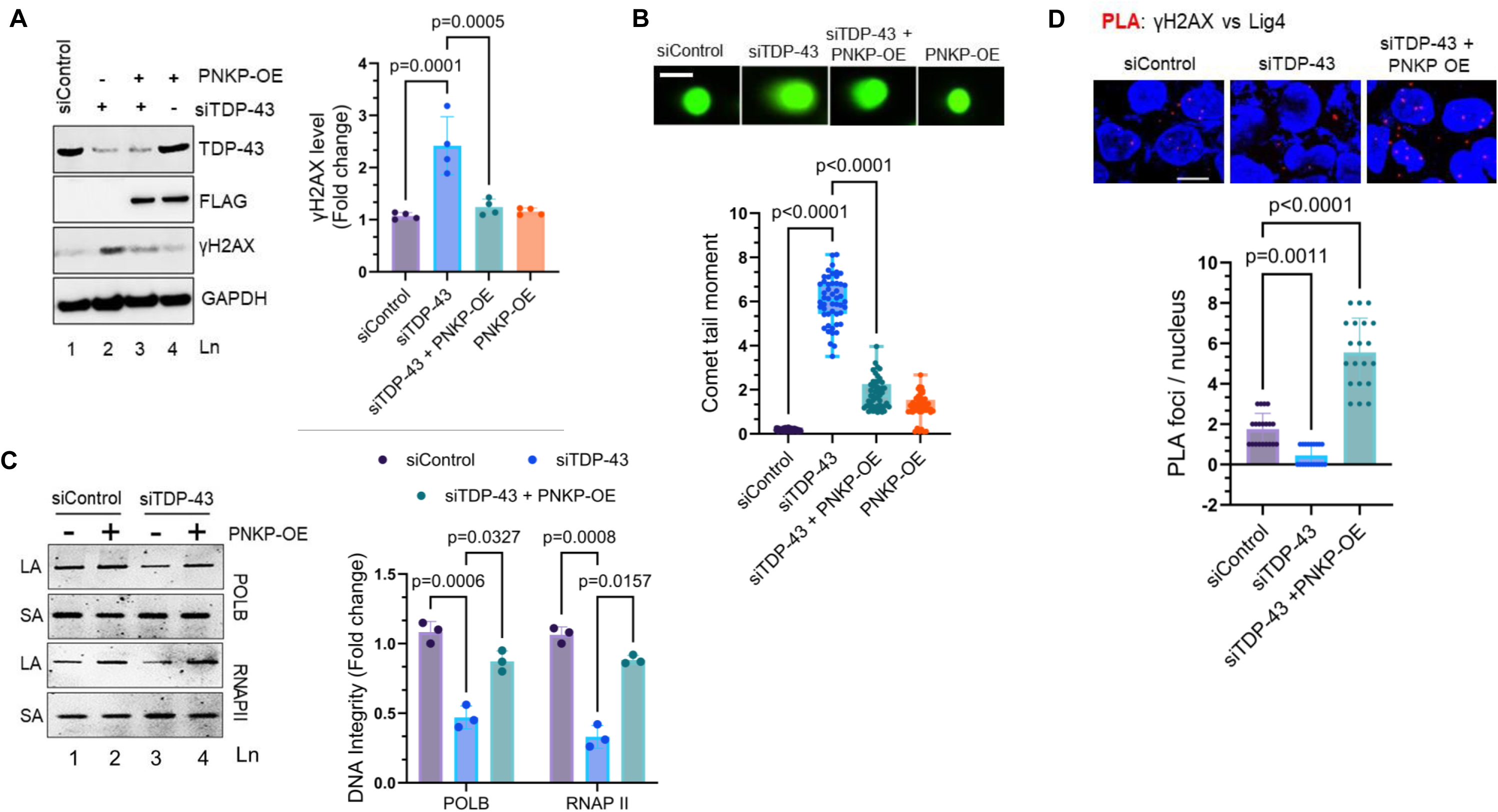
PNKP supplementation rescues DNA damage in TDP-43 KD cells. **A**. IB showing γH2AX levels in TDP-43 KD (siTDP-43) HEK293 cells with or without PNKP overexpression (OE). The histogram shows quantitation of γH2AX levels (band intensity, normalized to GAPDH). **B**. Neutral comet assay and quantitation of tail moment. N = 50 cells. Scale bar, 10 µm. **C**. DNA integrity analysis using LA-qPCR of POLB (12.4 kb) and RNAPII (11.3 kb) genomic regions. Quantitation of PCR products was performed using the Pico-Green method in a plate reader. The long amplicons were normalized to their respective short amplicons of the POLB (196 bp) and RNAPII (295 bp) genes and represented in the bar diagram as fold change of DNA integrity. **D**. PLA of Lig4 vs γH2AX shows reduced signal in TDP-43 KD HEK293 cells, which was rescued after PNKP-OE. Scale bar, 10 µm. The histogram shows the quantitation of the number of PLA foci per nucleus counted from 25 cells. The groups were compared using one-way or two-way ANOVA. Error bars represent mean ± SD; significance at p ≤ 0.05.

To further confirm the effects of PNKP complementation on DNA integrity, LA-qPCR was performed by amplifying 10-12 kb fragments encompassing two actively transcribed genes - DNA polymerase beta (POLB) and RNA polymerase II (RNAPII). The DNA damage analysis demonstrated a significant recovery of genome integrity as shown by increased PCR band intensity in PNKP-OE cells treated with siTDP-43 compared to cells treated with siTDP-43 alone. The long amplicon product levels were normalized with short amplicons from the same genes for DNA damage analysis (**Figure 3C**).

Our recent studies suggest that the efficient recruitment of XRCC4/DNA Lig4, the complex capable of covalently sealing the ends of a DSB, is compromised in the absence of functional nuclear TDP-43 (5, 8). However, the mechanism underlying this disruption remained elusive. Given PNKP’s critical role in processing damaged DNA termini at DSB sites to generate ligatable 3’-OH and 5’-P ends, we investigated whether the loss of PNKP activity contributes to the impaired recruitment of Lig4 complexes to the DSB sites. PLA revealed a significant reduction in Lig4 interactions with γH2AX-marked DSB sites in the nuclei of TDP-43 KD cells (**Figure 3D**). This deficiency was effectively rescued by PNKP-OE, suggesting that the restoration of PNKP activity may enhance the recruitment of DNA Lig4 to genomic DSBs.

### Disrupted TDP-43–PNKP association in the TC-NHEJ complex increased damage in transcribed genome in TDP-43 proteinopathy

Our previous studies uncovered the presence of a pre-formed TC-NHEJ complex involving PNKP and other DSB repair proteins in mammalian brain samples (31, 36). Notably, PNKP plays a crucial role in RNA-templated error-free TC-NHEJ repair, thereby protecting the transcribed genome (17, 31, 32). Since our results indicate that either depletion or mislocalization of TDP-43 is sufficient to disrupt the PNKP activity, we assessed PNKP’s association with the TC-NHEJ components in nuclear fractionates from autopsied cortical brain samples of ALS and age-matched control subjects using PNKP co-IP assay. Importantly, the interaction of PNKP with TDP-43 was significantly compromised in ALS, indicating potential involvement of TDP-43 in the TC-NHEJ pathway to facilitate PNKP-mediated DSB repair in the transcribed genomic regions (**Figure 4A)**. We also observed a concomitant decrease in the association of other TC-NHEJ proteins, such as Lig4 and PFKFB3, in ALS (**Figure 4A**). This result was consistent with our previous finding showing disruption of the C-NHEJ repair pathway in TDP-43 pathology.

**Figure 4.**
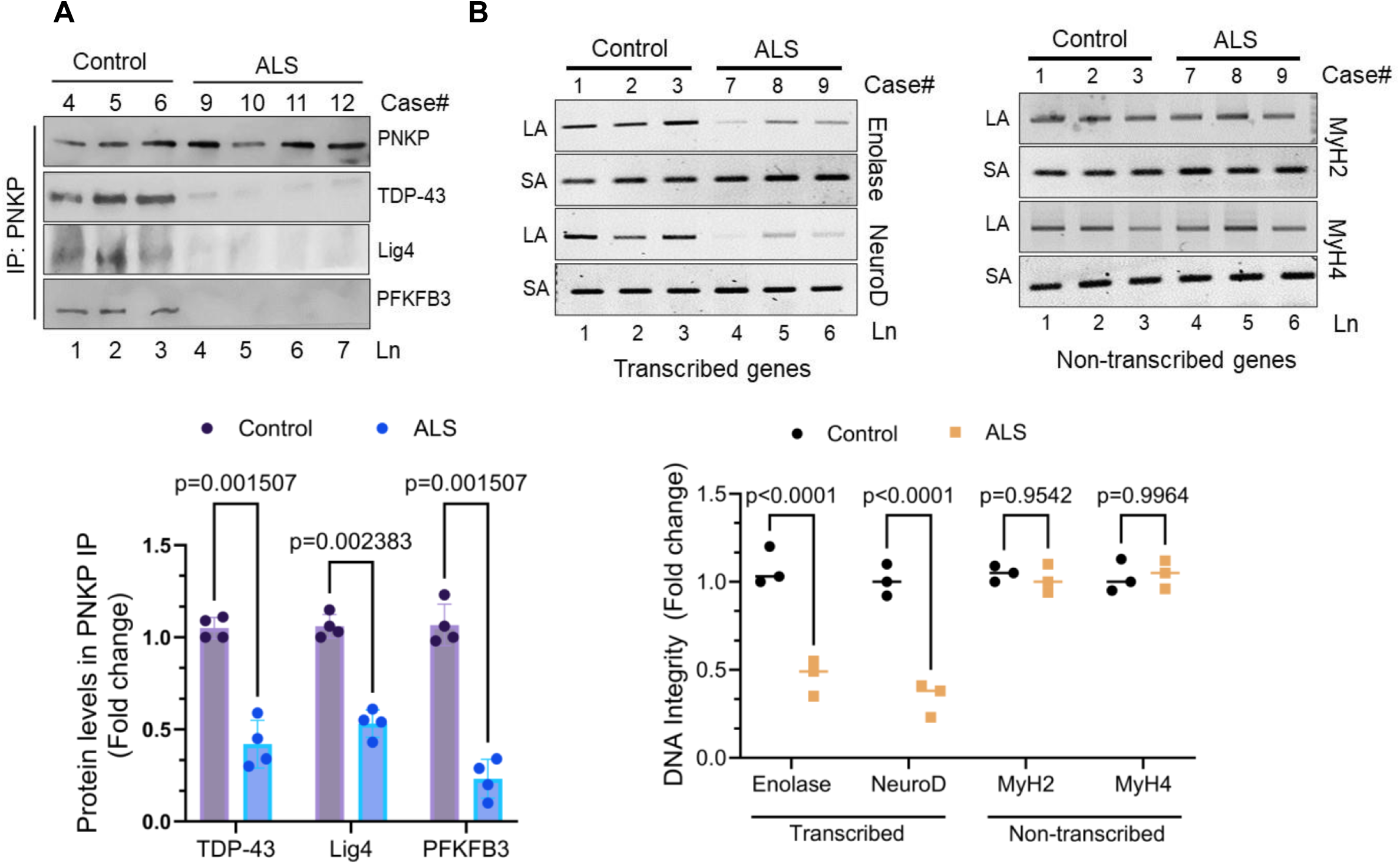
Partial characterization of the TC-NHEJ complex in post-mortem patient tissues and preferential accumulation of genome damage at transcribed genes in TDP-43 pathology. **A**. NE from autopsied frontal cortex tissues of human ALS patients (Lns 4-7) and non-neurological controls (Lns 1-3) (frontal cortex) were IP’d with anti-PNKP antibody and tested for associated proteins (as indicated). Quantitation of IP’d protein levels in terms of relative band intensity. **B**. Representative agarose gel images showing accumulation of genome damage in transcribed (Enolase & NeuroD) and non-transcribed (MyH2 & MyH4) genes in autopsied brain samples of non-neurological control and ALS-TDP-43 patients (N = 3) by LA-qPCR. The PCR products were independently quantified using the Pico-Green in a plate reader. Relative long amplicon values were normalized to their respective short amplicon ones: all LA amplicons were 6 kb in size and SA of all indicated genes were 300 bp, except for NeuroD, which was 200 bp. The histogram shows the fold change of DNA integrity. The groups were compared using two-tailed multiple t-tests. Error bars represent mean ± SD; significance at p ≤ 0.05.

Since PNKP is involved in the preferential repair of the transcribed genome, we assessed genome damage in actively transcribed versus non-transcribed regions in genomic DNA from cortical tissues of ALS patients versus age-matched control tissues. LA-qPCR analysis revealed a significant increase in DSB accumulation in transcribed genes (*left panel*; Enolase and NeuroD) compared to non-transcribed genes (*right panel*; MyH2 and MyH4) in ALS-TDP-43 tissues (**Figure 4B**). This preferential accumulation of DNA damage in transcriptionally active regions was consistent with a decrease in PNKP activity and impaired TC-NHEJ repair in ALS.

### Reduced PFKFB3 levels correlate with compromised PNKP activity and supplementation with F2,6BP restores PNKP activity

We have demonstrated that F2,6BP, the product of PFKFB3, is a positive co-regulator of PNKP activity *in vivo* (36). We further showed reduced levels of PFKFB3 and F2,6BP in the affected regions of HD and SCA3 patient brain samples, a plausible cause of PNKP inactivation in these pathologies (36). Therefore, to elucidate the mechanism of near-complete loss of PNKP activity in ALS/FTD, we assessed levels of PFKFB3 and F2,6BP in patients versus age-matched non-neurological controls. IB analysis revealed significant decreases in PFKFB3 levels in cortical samples from ALS and FTD patients compared to age-matched controls (**Figure 5A**; *Supplementary* **Figure S1B**). Our results corroborated earlier reports indicating a more than two-fold decrease in the PFKFB3 expression in the cortex of post-mortem ALS patients (51, 52). This result was also consistent with the observation of reduced PFKFB3 levels associated with the depletion of TDP-43 in HEK293 cells (**Figure 2A**). We further observed that the levels of F2,6BP were significantly lower in ALS/FTD patient brain samples compared to age-matched controls (**Figure 5B**), correlating with the reduced levels of PFKFB3 in those samples. These data further suggest a potential link between disrupted glycolytic regulation and impaired DNA repair mechanisms under neurodegenerative conditions. Next, we tested the effect of exogenous F2,6BP supplementation in restoring nuclear PNKP activity in patient samples. Notably, supplementation with F2,6BP significantly restored PNKP activity in the NE of ALS brain samples (**Figure 5C**; Lns 7-10 vs. Lns 3-6). Furthermore, we observed F2,6BP-mediated restoration of PNKP activity in a dose-dependent manner both in sporadic ALS (**Figure 5D**, Lns 5-6) and Guam-ALS samples **(Figure 5E**, Lns 5-7). Notably, we found that F2,6BP but not F1,6BP or F6P could restore PNKP activity in ALS/FTD, which was consistent with our previous finding in polyQ diseases (36). These results also highlight the importance of restoring the F2,6BP level to stabilize PNKP activity, even under stress conditions, thus indicating a metabolic reprogramming-mediated novel regulatory pathway for restoring DSB repair machinery and genome integrity in neurodegenerative diseases.

**Figure 5.**
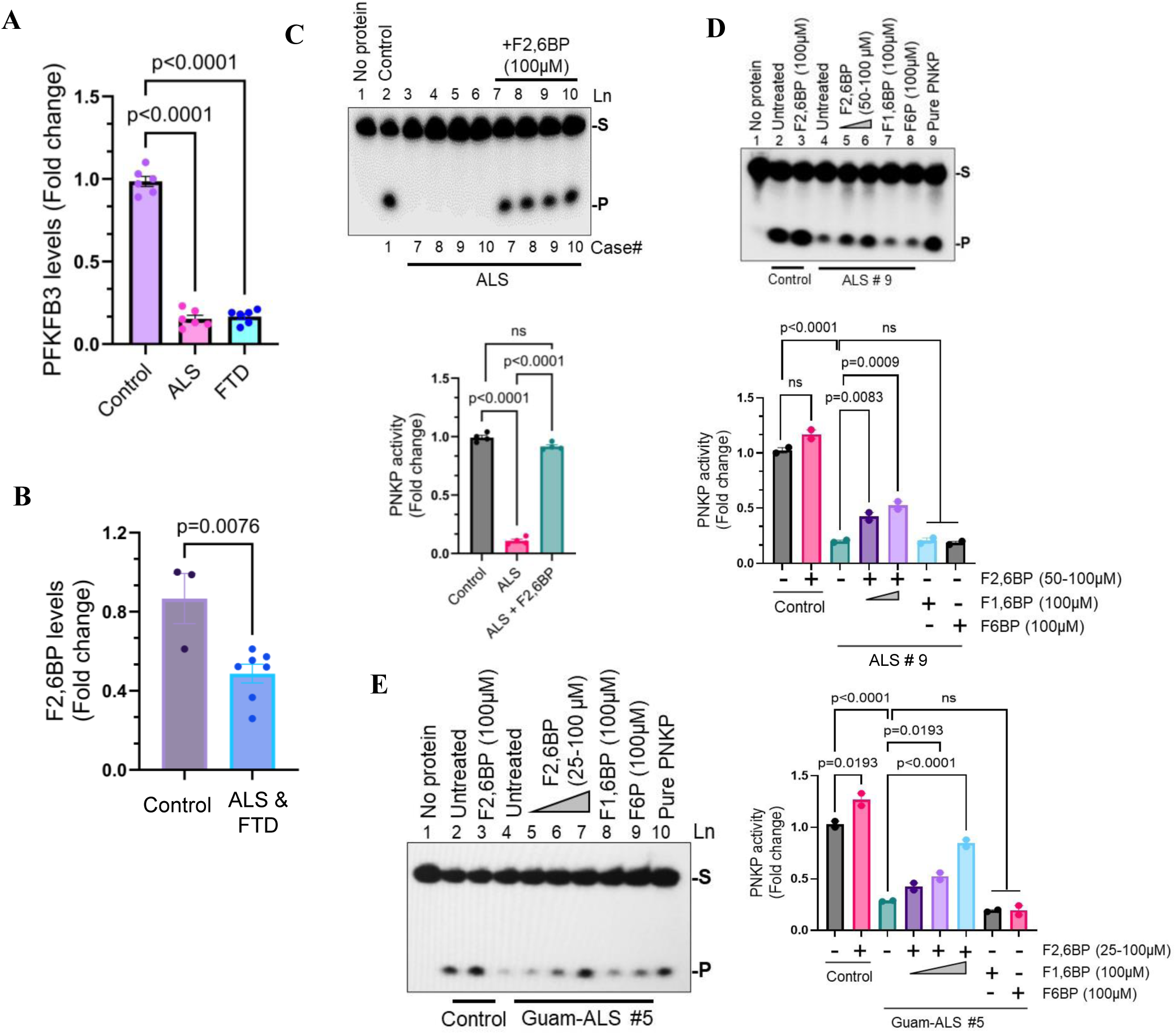
Reduced PFKFB3 and F2,6BP levels in ALS/FTD brains and restoration of PNKP activity by F2,6BP supplementation. **A)** The bar graph illustrates relative levels of PFKFB3 in ALS and FTD patients vs. non-neurological controls normalized with HDAC2. Data were analyzed using one-way ANOVA. **B)** Quantitation of F2,6BP levels in control (N=3), ALS (N=3) and FTD (N=4) patient frontal cortices. **C)** Representative image showing 3’-phosphatase activity of PNKP in NE from ALS-TDP-43 samples (Lns 3-6) compared to age-matched non-neurological controls (Ln 2), with activity restored upon adding the F2,6BP metabolite (Lns 7-10). Lane 1 shows the substrate-only negative control. The histogram quantifies PNKP activity as fold change, analyzed by one-way ANOVA. S: substrate, P: released phosphate. **D)** Representative image showing the concentration-dependent rescue of PNKP activity (Lns 5-6) by F2,6BP in a sporadic ALS-TDP-43 sample (Ln 4), compared with an age-matched control (Ln 2). F1,6BP and F6P were used as negative controls (Lns 7, 8). Lane 1 shows the substrate-only, lane 9: purified PNKP (25 fmol). The histogram presents relative PNKP activity as fold change, with analysis by two-way ANOVA. **E)** Representative image demonstrating concentration-dependent restoration of PNKP activity by F2,6BP in a Guam-ALS-TDP-43 sample compared to an age-matched control, with F1,6BP and F6P serving as negative controls. The histogram represents relative PNKP activities as fold change, analyzed via two-way ANOVA. Error bars represent mean ± SD; significance at p ≤ 0.05; ns = non-significant.

### F2,6BP supplementation restores PNKP activity in ALS patient-derived NPSC lines

To elucidate the impact of ALS and FTD-associated TDP-43 mutations on DNA repair mechanisms, we utilized three different ALS and one FTD patient iPSC-derived NPSC lines. Initially, we focused on two ALS-linked TDP-43^G287S^ and TDP-43^G298S^ mutations. **Figure 6A** illustrates the domain location of the G287S mutation in TDP-43 protein and a CRISPR/Cas9-mediated mutation-corrected isogenic TDP-43^G287G^ NPSC line (illustration in **Figure 6B**). To elucidate the mechanistic effect of these TDP-43 mutants on the PNKP activity and associated DSB repair proficiency, we first assessed the extent of cytosolic mislocalization of TDP-43^G287S^ in this patient-derived NPSC line. Compared to the isogenic control line, TDP-43^G287S^ cells exhibited a significantly higher nuclear-to-cytosolic ratio (N/C; P < 0.000001) of subcellular distribution of TDP-43 (**Figure 6C**). Furthermore, we also examined this phenomenon by IB analyses of insoluble fractionates from ALS patient-derived TDP-43^G287S^ NPSCs in comparison to its isogenic control TDP-43^G287G^ line, which showed significantly increased accumulations of pathological phenotypes of TDP-43 as well as increased levels of polyubiquitinated proteins in TDP-43 mutant line vs. the isogenic control (**Figure 6D**). Notably, treatment with F2,6BP (100 µM, 72 h) significantly reduced such pathogenic protein aggregates and polyubiquitination level. Based on these characterizations, we next examined PFKFB3 levels and 3’-phosphatase activity of PNKP in the TDP-43^G287S^ mutant vs. its isogenic control to correlate with our earlier results in ALS/FTD patient samples. IB analysis revealed a significant reduction (∼60%) in the PFKFB3 protein level in TDP-43^G287S^ NPSCs relative to their isogenic control cells, while PNKP protein levels remained unaffected (**Figure 6E**). A marked decrease in PNKP activity correlated with reduced PFKFB3 levels in these samples, and this loss of activity was restored in a dose-dependent manner by F2,6BP supplementation *in vitro* (**Figure 6F**).

**Figure 6.**
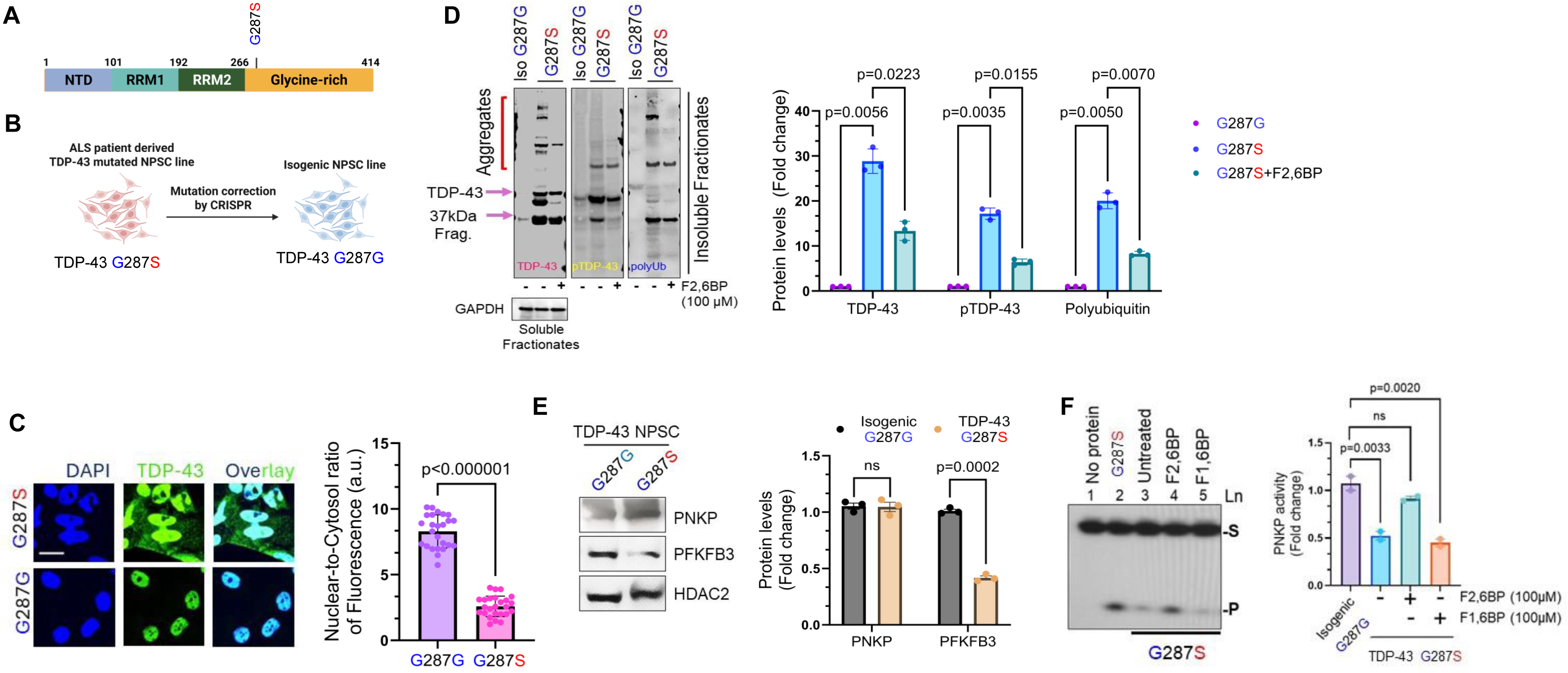
F2,6BP rescues DNA damage caused by mutant TDP-43–mediated PNKP inhibition in ALS-TDP-43G287S patient-derived NPSC lines. **A)** Schematic representation of TDP-43 protein domains, indicating the G287S mutation site. **B)** An illustration of the CRISPR/Cas9-mediated correction of the TDP-43G287S mutation to generate an isogenic TDP-43G287G iPSC line. **C)** IF analysis showing cytosolic mislocalization of TDP-43 in the mutant cells. Quantitation of nuclear-to-cytosol ratio of TDP-43 fluorescence (in arbitrary units, a.u.). The groups were compared by t-test. Scale bar, 10µm. **D)** IB analysis of insoluble fractionates from the isogenic and TDP-43G287S cells with or without F2,6BP (100 µM) treatment for total TDP-43, pTDP-43 (S409/S410) and polyubiquitinated proteins. The GAPDH served as the loading control indicating the uniformity in preparing insoluble fractionates. Bar graphs show changes in protein levels of the indicated target proteins as fold changes. **E)** IB analysis of PNKP and PFKFB3 levels in the NE from G287S mutant and isogenic G287G NPSCs. HDAC2 served as the loading control. Quantification of levels of PFKFB3 and PNKP between the groups is shown on the right. **F)** Representative gel image displaying PNKP activities in NE from the mutant (Ln 3) and isogenic NPSCs (Ln 2) and its rescue by F2,6BP (100 µM, Ln 4) but not F1,6BP (Ln 5). Bar diagram shows quantification of the PNKP activity between the groups (N=3). S: substrate, P: released phosphate. Lane 1, no protein. The groups were compared using t-test, one-way or two-way ANOVA as appropriate. Error bars represent mean ± SD; significance at p ≤ 0.05; ns = non-significant.

We next extended our analysis to the ALS patient-derived TDP-43^G298S^ mutant NPSCs. Similar to the TDP-43^G287S^ mutant, IF analyses showed a significantly enhanced cytosolic mislocalization of TDP-43 with ∼ four-fold less N/C ratio (P < 0.0001) (*Supplementary* **Figure S2A**). IB analysis revealed a substantial elevation in γH2AX level in the G298S mutant, indicating elevated DNA damage, while total H2AII levels remained comparable between mutant and WT cells (*Supplementary* **Figure S2B**). Finally, we assessed PNKP activity in the G298S mutant cells. Similar to our findings with the G287S mutation, we observed an ∼80% reduction in PNKP activity in TDP-43^G287S^ NPSCs compared to WT control cells, while PNKP activity could be restored by F2,6BP in a dose-dependent manner (*Supplementary* **Figure S2C**).

Further investigations extended to an engineered ALS-linked TDP-43^Q331K^ mutant, and its isogenic control (*Supplementary* **Figure S3A**) revealed similar patterns of TDP-43 mislocalization (*Supplementary* **Figure S3B**), reduced PFKFB3 levels (*Supplementary* **Figure S3C**), and decreased PNKP activity (*Supplementary* **Figure S3D**), compared to respective controls. However, PNKP activity was restored by F2,6BP supplementation **(Figure S3D)**. In addition, we examined an FTD patient-derived cell line harboring both TDP-43^A382T^ and a C9ORF72 repeat expansion (*Supplementary* **Figure S3E**). The results demonstrated a decrease in PNKP activity in mutant iPSCs and restoration of PNKP activity upon F2,6BP supplementation, which was consistent with other TDP-43 mutant cell lines (*Supplementary* **Figure S3F**), underscoring the potential of F2,6BP in restoring DNA repair capacity in TDP-43 pathology. Collectively, our data demonstrates that reduced PNKP activity is linked to decreased levels of PFKFB3 and its product F2,6BP, a crucial cofactor for PNKP activity, under pathological conditions.

### Altered PNKP activity and its restoration by F2,6BP in ALS-TDP-43 mouse brain extracts

Next, we investigated the impact of TDP-43 pathology on PNKP level and its activity in an ALS-TDP-43 mouse model. **Figure 7A** shows the schematic of the genetic construct employed to drive murine Tdp-43^ΔNLS^ mutant expression. We have shown that murine Tdp-43^ΔNLS^ expression can enhance the pathological phosphorylation and simultaneous cytosolic aggregation of endogenous Tdp-43, leading to genome instability and inflammation (9).

**Figure 7.**
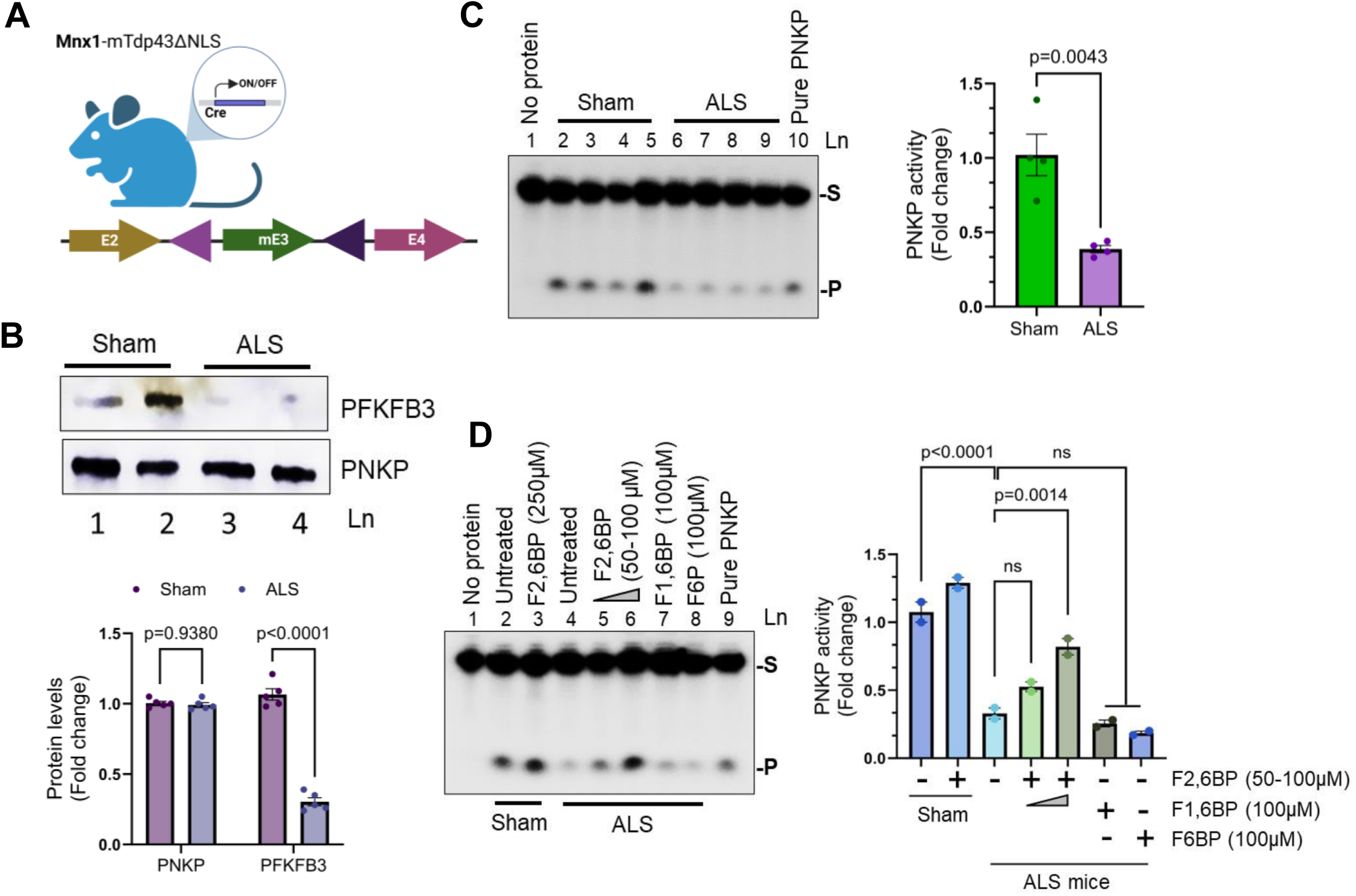
Reduced PNKP activity in TDP-43 mouse brain tissues correlates with PFKFB3 levels and its restoration by F2,6BP. Schematic of the construct used to generate Tdp43-ΔNLS expression under the UBC promoter in C57BL6 mice. **B**) Representative IB images showing PNKP and PFKFB3 levels in sham and ALS cortical samples. Bar graph displaying changes in PNKP and PFKFB3 levels across the groups as fold changes. Data are analyzed by one-way ANOVA. **C)** Representative image demonstrating reduced PNKP activity in the nuclear extracts of ALS (Lns 6-9) compared to age-matched sham controls (Lns 2-5). Ln 1: substrate only (negative control); Ln 10: substrate plus pure PNKP protein as positive control. The histogram displays the quantification of PNKP activity as fold change, analyzed by Student’s t-test. S: substrate, P: released phosphate. **D)** Representative image showing the rescue of PNKP activity by F2,6BP in a dose-dependent manner in ALS mice sample compared to WT, where F1,6BP and F6P served as negative controls. Histogram indicating relative PNKP activities as fold change across the groups. Data were analyzed by two-way ANOVA. Error bars represent mean ± SD; significance at p ≤ 0.05; ns = non-significant.

We next assessed the levels of PFKFB3 and PNKP in cortical samples from ALS and sham (wild type) mice. IB analysis revealed a significant reduction in PFKFB3 protein levels in the NE of ALS mice compared to controls, while PNKP levels remained unaltered (**Figure 7B**). We further evaluated 3 -phosphatase activity of PNKP in the NE from cortical tissues of ALS and age-matched sham mice (**Figure 7C**), which displayed markedly decreased PNKP activity in ALS mouse brains (Lns 6-9) relative to sham controls (Lns 2-5). Importantly, we observed a dose-dependent increase in PNKP activity following F2,6BP supplementation, whereas the glycolytic metabolites F1,6BP and F6P did not produce similar effects (**Figure 7D**). Overall, these results indicate that TDP-43 pathology in ALS mice leads to significant reductions in PNKP activity. This effect correlated with reduced PFKFB3 levels, which was supported by our results that treatment with F2,6BP could effectively rescue PNKP function, suggesting a promising metabolic approach to restoring DNA repair in ALS animal models.

### Supplementation of F2,6BP restored genome integrity and partially rescued motor phenotype in a *Drosophila* model of ALS

To test the effect of F2,6BP in reversing neurodegenerative phenotype in ALS, we used a fruit fly (*Drosophila melanogaster*) model of ALS where humanized TDP-43^Q331K^ mutant was expressed under UAS (BDSC# 79590) and crossed with the driver line (BDSC# 64199) to drive motor neuron-specific expression of the mutant TDP-43, a pathogenic variant known to cause early synaptic dysfunction and subsequent motor deficits (53). The flies were then treated with 1-2 µl of either 50 µM F2,6BP or mock buffer (as control) for 21 days and the motor phenotype was tested by climbing assays, which is a robust and sensitive measure of locomotor function that reliably declines with age and neurodegeneration in fly models of disease (54). We observed a significant increase in climbing score after supplementation with F2,6BP compared to mock buffer control (**Figure 8A**). We further examined the DNA strand break accumulation in *Drosophila CrebB* and *Neurexin* genes by LA-qPCR. These genes were selected not only for their critical roles in synaptic function, learning, and memory (55, 56) but also because their large size makes them particularly susceptible to transcription-associated DNA damage, a known vulnerability in neurodegenerative conditions (57). We observed an elevated level of DNA damage following Q331K expression compared to the flies used to generate the strains (w1118) (**Figure 8B**, Ln 2 vs. Ln 1). These data further indicates that the mock treatment failed to restore genome integrity. However, significant repair was observed following F2,6BP supplementation (**Figure 8B**, Ln 3 vs. Ln 2), and the genome integrity was comparable to that of w1118, which correlated with the rescue of motor phenotype. These findings confirmed the potential of F2,6BP to specifically reverse the ALS neurodegenerative phenotype in a living organism.

**Figure 8.**
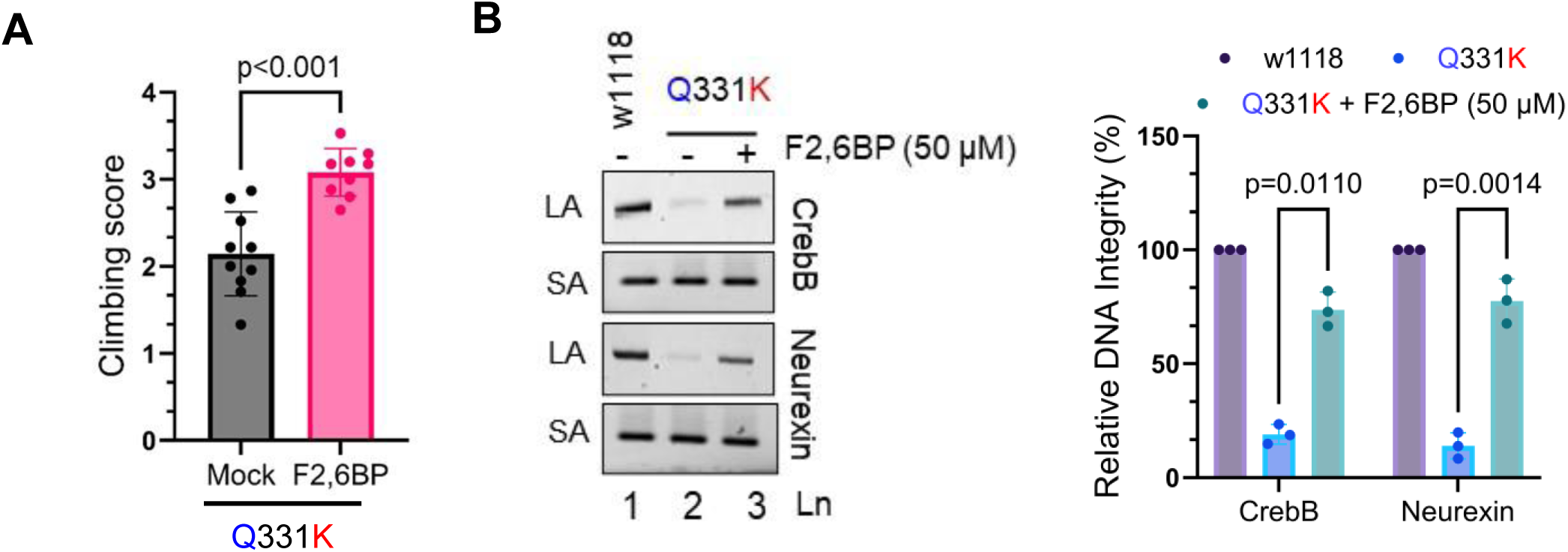
F2,6BP rescued motor deficiency and restored genome repair in a whole organism. **A**) Climbing score for ALS flies expressing TDP-43 gene with disease-linked mutation Q331K in motor neurons (10 flies in each cohort, male and female mixed) treated with either mock buffer or F2,6BP (50 µM). One-way ANOVA was used to assess the statistical significance. **B**) Representative agarose gel images of long (∼8 kb; LA) and short (∼200 bp; SA) amplicons of the *CrebB* and *Neurexin* genes from flies either mock-treated (lane 2) or treated with F2,6BP (Ln 3). Lane 1: w1118. **C**) The normalized relative band intensities with w1118 were arbitrarily set as 100 (N = 3). Error bars represent mean ± SD; significance at p ≤ 0.05.

## Discussion

The discovery of TDP-43 pathology, initially identified due to its cytosolic mislocalization in motor neurons of ALS and FTD patients (1, 2, 58–60), has profoundly shaped our understanding of neurodegenerative diseases (61, 62). Beyond ALS and FTD, TDP-43 pathology is now recognized in multiple forms of dementia, including Alzheimer’s disease (AD) and related dementias (ADRD), where over 50% of patients exhibit TDP-43 pathologies (63, 64). This broader involvement highlights the importance of dissecting the mechanistic underpinnings of TDP-43-driven cellular dysfunction.

We and others have shown that PNKP plays a critical role in DNA DSB repair through a C-NHEJ pathway (20, 65). Our current findings reveal that TDP-43 mislocalization in ALS and FTD results in a severe loss of 3’-phosphatase activity of PNKP, essential for generating ligatable DNA ends, without affecting its protein levels. This defect is tightly associated with reduced levels of glycolytic enzyme, PFKFB3, and its metabolic product, F2,6BP, underscoring a novel metabolic regulation of genome repair. We have recently shown that PNKP can bind F2,6BP with high affinity (Kd=0.5 µM) (42) and thus can utilize F2,6BP as a cofactor for its activity, which further supports our observations. Importantly, supplementation with F2,6BP rescued PNKP activity in ALS/FTD brain extracts, patient iPSC-derived NPSCs, and an ALS mouse model, restoring genome integrity in transcribed regions.

This study expands the mechanistic link between metabolic dysregulation and DNA repair deficiency in neurodegeneration. Our prior reports on polyglutamine disorders such as HD and SCA3 had already implicated PNKP inactivation as an early event in neuronal dysfunction (31, 36, 65). Our findings extend this paradigm to ALS and FTD, unravelling a common mechanism of pathogenicity in various neurodegenerative diseases, wherein impaired PNKP activity, due to reduced F2,6BP, leads to accumulation of persistent DNA breaks and neurotoxicity, as shown schematically in **Figure 9**.

**Figure 9.**
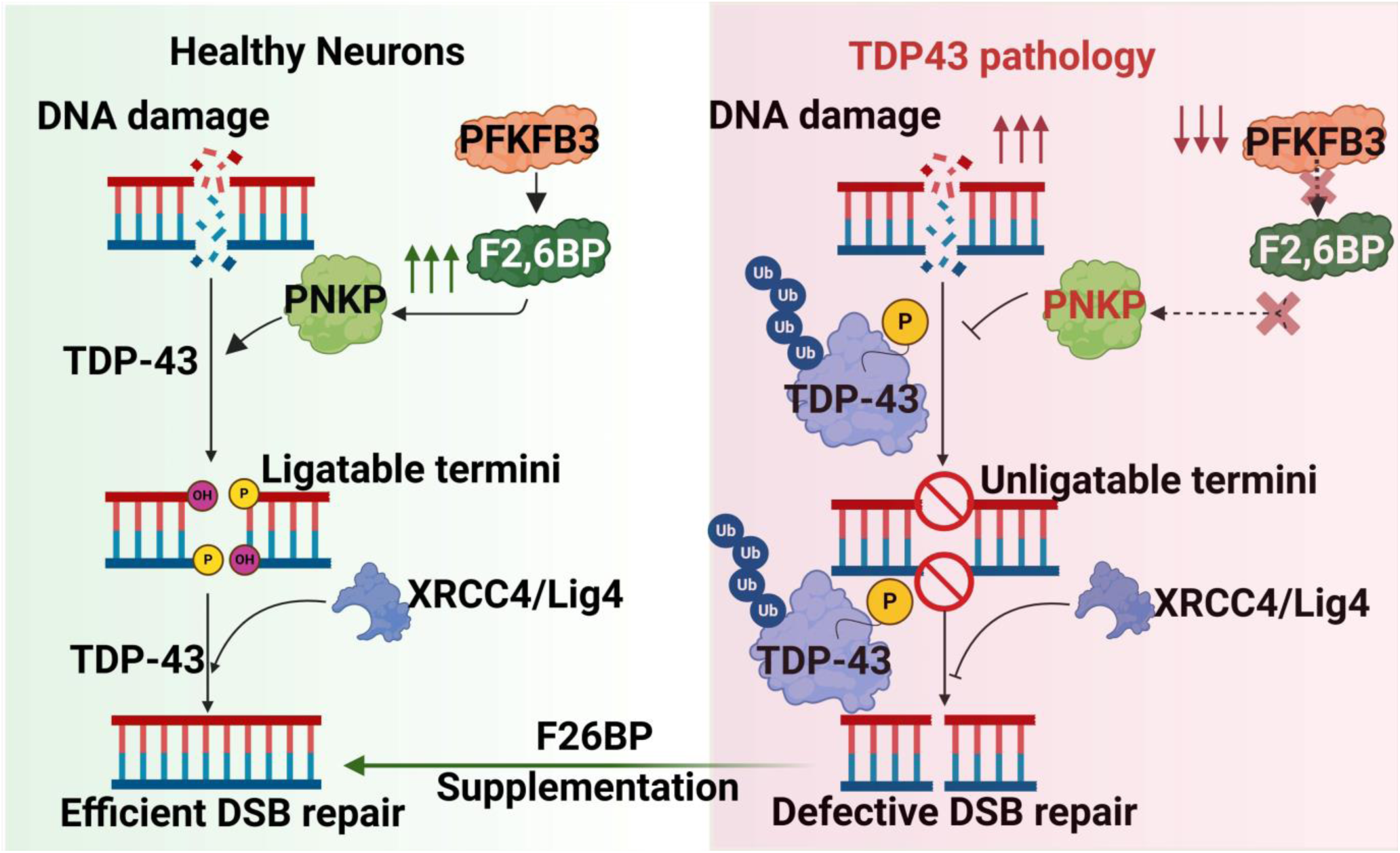
A Schematic Overview of PNKP Inhibition in Neurodegenerative Diseases with TDP-43 Pathology. The glycolytic enzyme PFKFB3 and its product, the small molecule metabolite F2,6BP, plays a pivotal role in regulating PNKP activity and in the repair of genome damage via transcription-coupled NHEJ. TDP-43 pathology (hyper-ubiquitination and phosphorylation) results in reduced levels of PFKFB3/F2,6BP and significant loss of PNKP activity, which results in neuronal loss. Supplementation of F2,6BP restores PNKP activity, suggesting its therapeutic potential.

Importantly, in addition to restoring DNA repair, F2,6BP significantly reduced pathogenic TDP-43 aggregates, including polyubiquitinated and phosphorylated forms, in patient-derived cell lines. This dual effect highlights the multifunctional therapeutic potential of F2,6BP, not only as a metabolic cofactor rescuing DNA repair but also as a suppressor of TDP-43 aggregation and cytoplasmic mislocalization. Whether this modulation occurs through direct effects on protein homeostasis, redox balance, or the proteasomal pathway remains to be elucidated.

Our study also demonstrates that the TC-NHEJ complex is disrupted in ALS-TDP-43 conditions. Preferential accumulation of DSBs in actively transcribed genes observed in autopsied human brain tissues underscores the vulnerability of transcriptionally active chromatin to repair failure. Restoration of PNKP activity reinstated the recruitment of Lig4 to DSBs and reduced γH2AX signals, validating functional rescue of repair machinery.

A significant strength of this study lies in its cross-validation across multiple models, including sporadic ALS, Guam-ALS, and FTD patient autopsy tissue from two biorepositories, patient iPSC-derived neurons harboring mis-localizing mutations in TDP-43, a conditional ALS-Tdp-43 mouse model, and a *Drosophila* ALS model. The findings in the *Drosophil*a ALS model are particularly compelling, as they demonstrate a clear therapeutic effect in a whole, living organism. Importantly, F2,6BP supplementation not only corrected molecular defects but also improved motor phenotypes in flies, demonstrating translational potential. This tight correlation between the restoration of genome integrity and the improvement of the motor phenotype provides powerful *in vivo* evidence for the PNKP-PFKFB3-F2,6BP axis as a key therapeutic target. Furthermore, these results align with a growing body of literature implicating metabolic dysregulation as a central node in the pathophysiology of various neurodegenerative diseases (66).

One limitation of this study is the potential for F2,6BP to enhance glycolysis globally, which may have unintended consequences in non-neuronal tissues or exacerbate hypermetabolic conditions. Future studies should focus on optimizing the dose, delivery, and cell-type-specific targeting of F2,6BP or its analogs that retain DNA repair modulating activity without metabolic stimulation. Additionally, the mechanistic exploration of how TDP-43 pathology drives PFKFB3 degradation could uncover potential targets for disease reversal upstream of DNA repair failure.

In summary, our work identifies a previously unrecognized PNKP-PFKFB3-F2,6BP axis disrupted in TDP-43 proteinopathies. We demonstrate that metabolic repletion with F2,6BP restores genome integrity via rescuing PNKP activity and reduces TDP-43 pathology. These findings provide a compelling rationale for developing mechanism-guided therapeutic strategies targeting genome maintenance in ALS, FTD, and potentially other TDP-43-related dementias.

## Supplementary Information

The online version contains supplementary materials, which include *Supplementary* **Figures S1-3** and **Tables S1-2**.

## Acknowledgments

The authors thank Dr. Anna Dodson (Center for Neuroregeneration, Houston Methodist Research Institute) for scientific editing of the manuscript. The authors thank Drs. Van Damme and Van Den Bosch (University of Leuven, Belgium) for their generous supports with ALS patient-derived iPSC lines. We are also thankful to Melissa Nijs (Laboratory for Neurobiology, VIB-KU Leuven) for arranging and shipping patient cell lines. Autopsied ALS and non-neurological control specimens were provided by the Department of Veterans Affairs Biorepository: VA Merit review BX002466. The authors also acknowledge the invaluable contribution of the Binghamton Brain Biobank for providing Guamanian ALS-PD autopsied patient samples.

## Author contributions

M.L.H. and T.H. conceived and designed the project and edited the final manuscript. A.C. and J.M. performed most of the experiments and wrote the manuscript; V.H.M.R., M.K., S.M.M., S.K.G., S.G.S., V.V., and M.M. performed various experiments and analyzed data. R.M.G and I.R. provided ALS and FTD autopsied human patient samples as well as critical comments and suggestions for the study; L.V.D.B. provided ALS-TDP-43 patient-derived iPSC lines. B.K. supervised the *Drosophila* study and provided analytical thoughts and suggestions for ALS fly model experiments and data analysis and interpretation. G.G. supervised the F2,6BP synthesis, assay and critical analysis of data related to this assay. All authors participated in data analyses, interpretation and approved the submission of the manuscript for publication.

## Data availability

All relevant data generated and analyzed in this study are available in this manuscript, online supplementary information or upon reasonable request.

## Ethics Approval

The animal study protocol was approved by Houston Methodist’s Institutional Animal Care and Use Committee under the approval number ISO00008727.

