## Supplementary material for "Fructose-2,6-bisphosphate restores TDP-43 pathology-driven genome repair deficiency in motor neuron diseases"

Supplementary Material associated with this article includes three figures and two tables

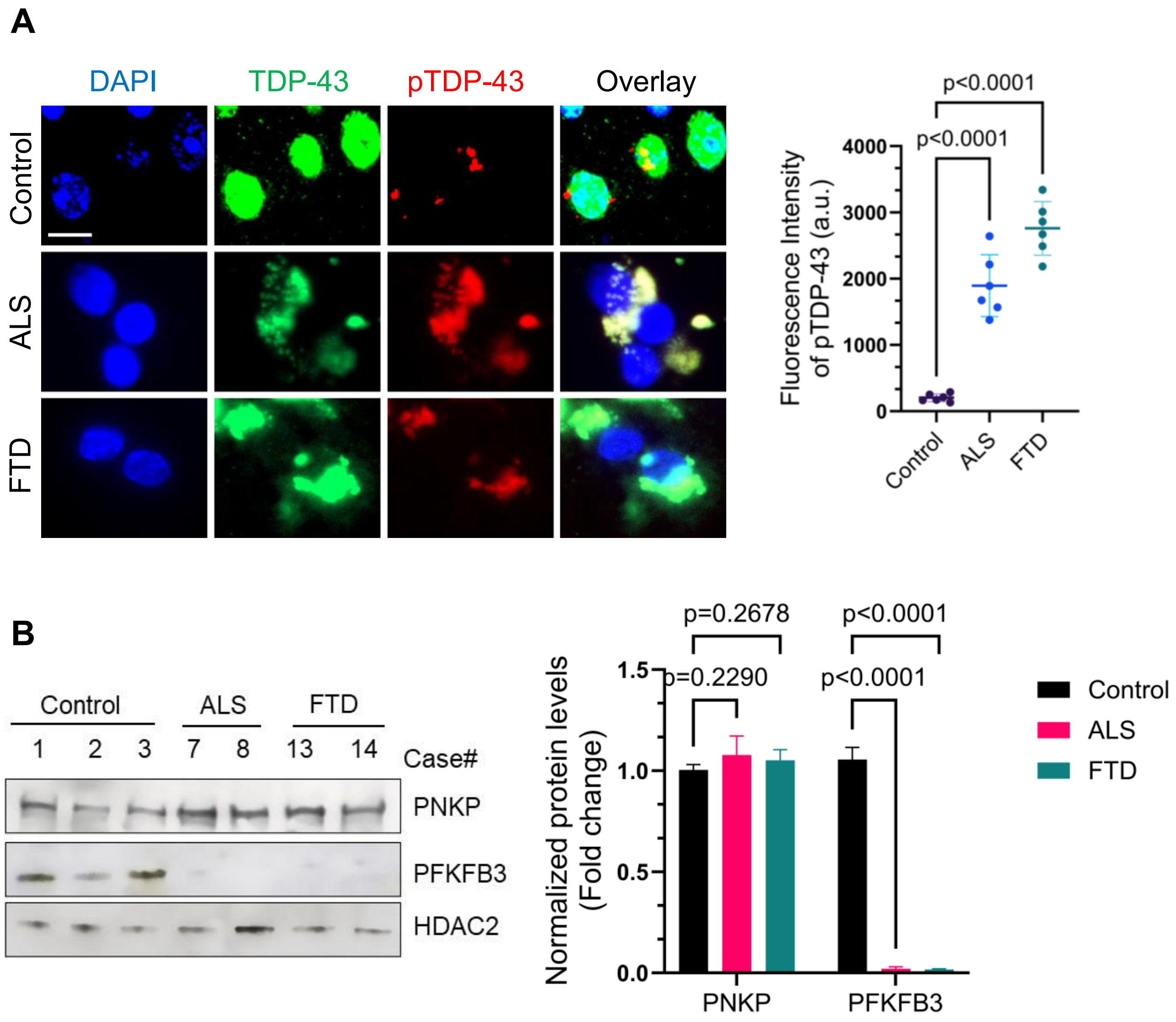


**Supplementary Figure S1 (Related to Figure 1).**

**A)** Representative immunofluorescence (IF) images showing cytosolic mislocalization of total TDP-43 and corresponding phosphorylated TDP-43 (pTDP-43, S409/410) expression levels in autopsied cortical sections from ALS, FTD, and age-matched non-neurological control samples (N = 6 cases per group). Scale bar, 10 μm. Quantification of fluorescence intensity (arbitrary units; a.u.) of pTDP-43 levels is included.

**B)** Representative IB image exhibiting PNKP and PFKFB3 levels in the nuclear extracts (NE) of cortical tissues from controls, ALS, and FTD patients, with HDAC2 serving as a loading control. Quantification of protein levels of PNKP and PFKFB3 across the study groups.

The groups were compared using the two-way ANOVA. Error bars represent mean ± SD; significance at $p \leq 0.05$; ns = non-significant.

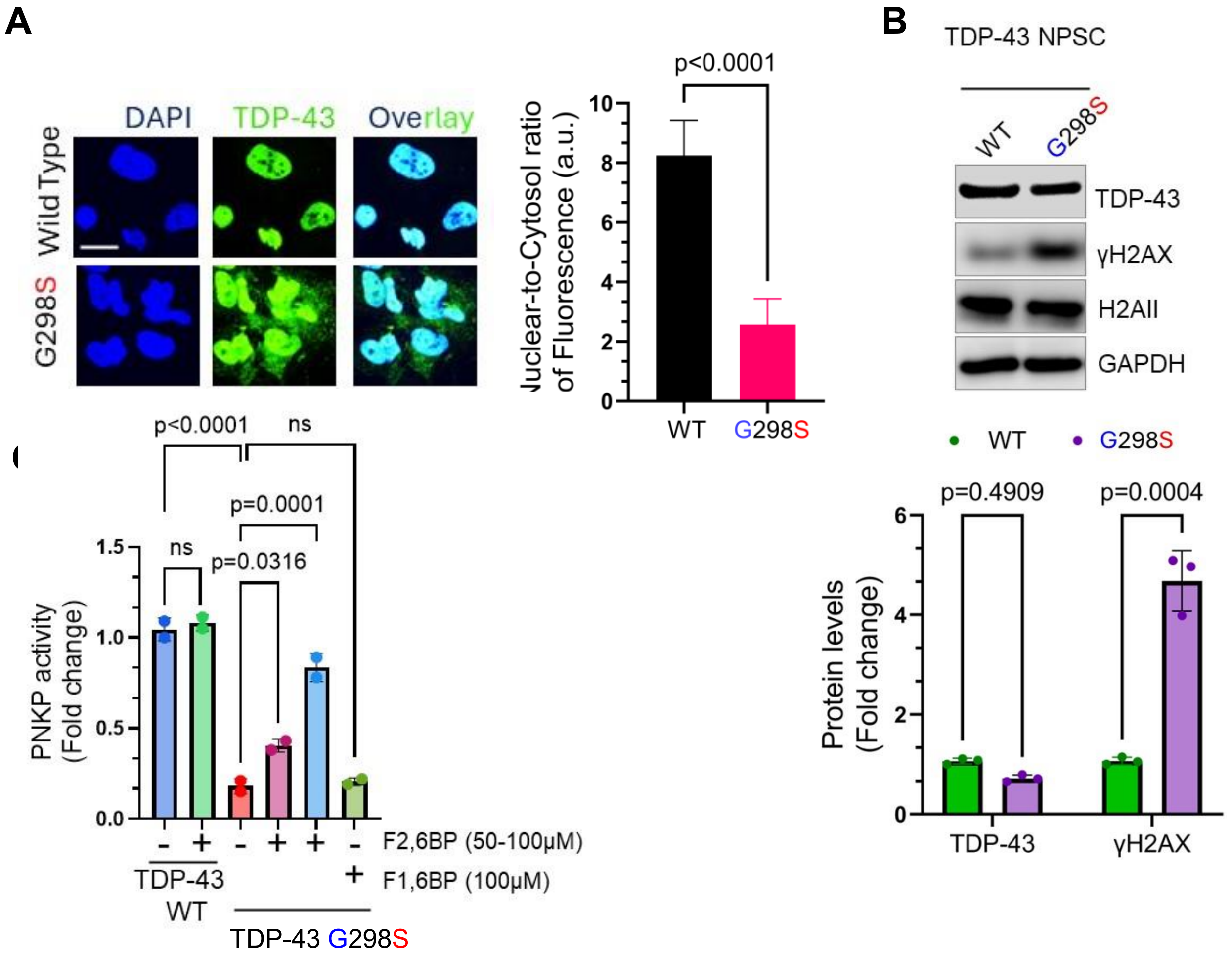


**Supplementary Figure S2 (Related Figure 6). F2,6BP rescues DNA damage caused by mutant TDP-43–mediated PNKP inhibition in an ALS-TDP-43$^{G298S}$ patient-derived NPSC lines.**

**A)** IF analysis of TDP-43's subcellular distributions in TDP-43 mutant versus wild-type (WT) control cells. Scale bar, 10 μm. Quantitation of nuclear-to-cytosol ratio of TDP-43 fluorescence signal intensities (a.u.).

**B**) IB analysis of γH2AX levels in TDP-43$^{WT}$ versus TDP-43$^{G298S}$ mutant cell extracts, where H2AII served as the loading control for γH2AX and GAPDH for whole cell extracts. Lower panel shows quantification of protein levels in fold change from three independent experiments is shown in the histogram.

**C**) Quantitation of PNKP activity in NE from TDP-43$^{WT}$ and TDP-43$^{G298S}$ mutant cell lines, with or without F2,6BP treatments (50-100 μM).

Error bars represent mean ± SD; the groups were compared for statistical significance using two-tailed t-tests or two-way ANOVA as appropriate; significance at $p \leq 0.05$; ns = non-significant.

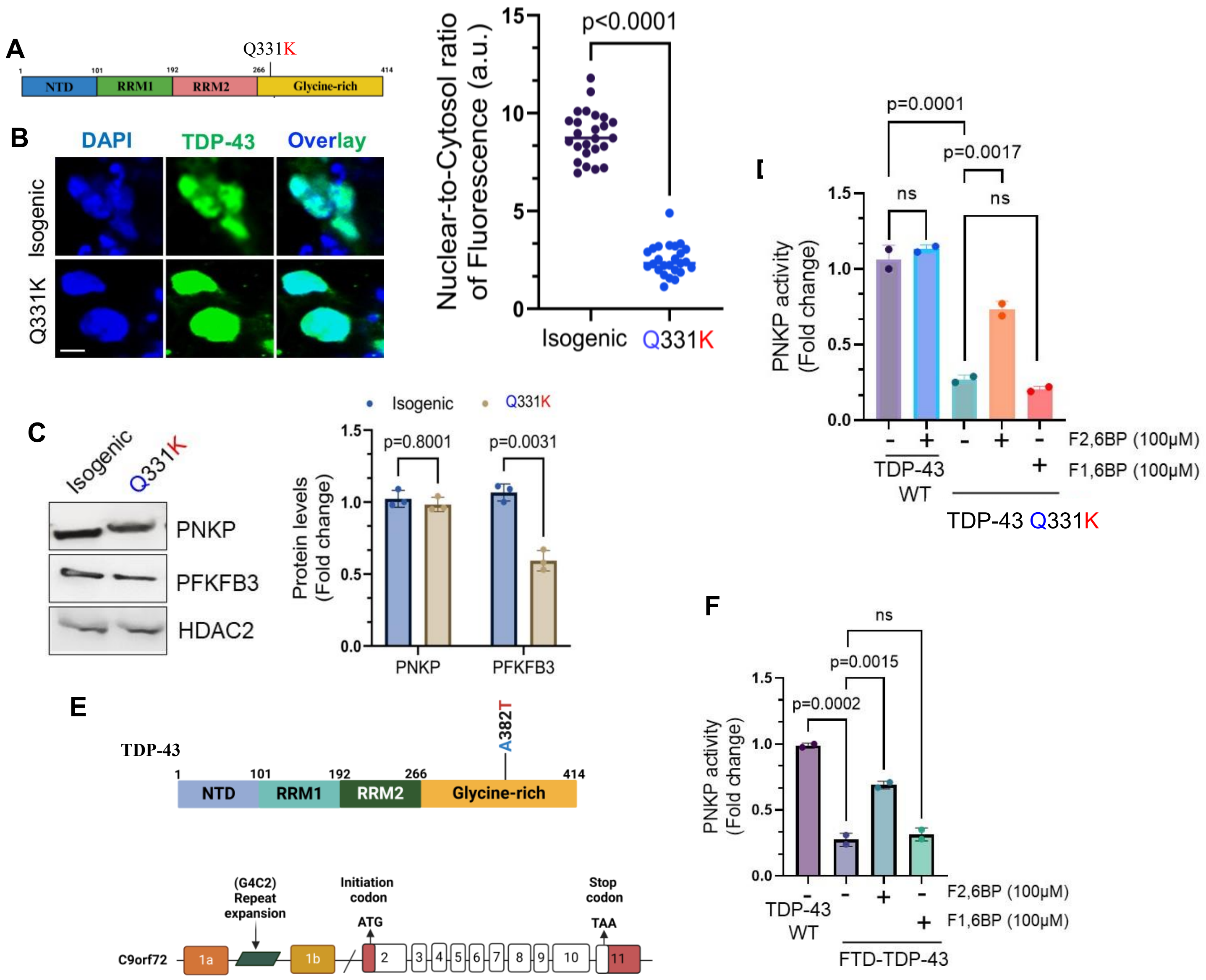


**Supplementary Figure S3 (Related Figure 6). DNA Damage Induced by Mutant TDP-43 via PNKP Inhibition is Rescued by F2,6BP in ALS Patient Cell Lines.**

**A**) Schematic representation of TDP-43 protein domains indicating the Q331K mutation site.

**B**) IF analysis of TDP-43's subcellular distributions in TDP-43 mutant versus wild-type (WT) control cells. Scale bar, 10 μm. Quantitation of nuclear-to-cytosol ratio of TDP-43 fluorescence signal intensities (a.u.).

**C**) IB analysis of PNKP and PFKFB3 levels in the NE from TDP-43$^{Isogenic}$ and TDP-43$^{Q331K}$ NPSCs. HDAC2 served as the loading control. The bar diagram shows the quantification of levels of PFKFB3 and PNKP between the groups.

**D**) Quantitation of PNKP activity in NE from TDP-43$^{Isogenic}$ and TDP-43$^{Q331K}$ mutant cell lines, with or without F2,6BP treatments (100 μM).

**E**) Schematic representation of an FTD patient-derived cell line harboring both TDP-43$^{A382T}$ and a C9ORF72 repeat mutation.

**F**) Quantitation of PNKP activity in NE from the FTD patient cell line, with or without F2,6BP (100 μM). F1,6BP (100 μM) treatment group served as the negative control. Error bars represent mean ± SD; the groups were compared for statistical significance using two-tailed t-tests or two-way ANOVA as appropriate; significance at $p \leq 0.05$; ns = non-significant.

**Supplementary Table S1:** Demographic and clinical characteristics of ALS and FTD-TDP-43 patients and age-matched non-neurological control subjects. The TDP-43 inclusion status, provided by the sourcing biorepositories and independently validated by us, is indicated.

| ID# | Age | Ethnicity | Sex | Pathology | Disease | TDP-43 pathology? |
|---|---|---|---|---|---|---|
| **Cohort 1** | | | | | | |
| 1 | 66 | Hisp/Latino | M | Bronchopneumonia | Non-neurological | No |
| 2 | 81 | White | F | Diffuse Alveolar Damage; Sepsis; acute Colitis | Selective neuronal necrosis | No |
| 3 | 62 | Hisp/Latino | M | Cardiopulmonary dysfunction; Hypertensive cardiomyopathy | Non-neurological | No |
| 4 | 68 | Whie | M | Bronchopneumonia and pleuritis; T-cell lymphoma; Renal dysfunction. | Non-neurological | No |
| 5 | 71 | White | F | Myocardial Ischemia; Aortic stenosis. | Non-neurological | No |
| 6 | 82 | Hisp/Latino | M | Cardiac ischemia | Non-neurological | No |
| 7 | 76 | White | M | Respiratory insufficiency; TDP-43 Inclusions; NFT; Braak II; Spinal degeneration. | Rapid onset ALS | Yes |
| 8 | 62 | African/Am | M | Spinal degeneration; Arteriosclerosis; TDP-43 Inclusions | ALS & Heart failure | Yes |
| 9 | 50 | White | M | Spinal degeneration; Midbrain degeneration; TDP-43 skein-like inclusions. | ALS & Respiratory failure | Yes |
| 10 | 80 | White | M | Spinal degeneration; NFT; Lewy body; Arteriosclerosis; Braak IV/VI. | ALS & Intercostal Respiratory failure | Yes |
| 11 | 58 | White | F | Spinal degeneration; Lewy bodies in olfactory bulb; vascular disease. | ALS | Yes |
| 12 | 83 | African/Am | M | Spinal degeneration; Braak III/VI; NFT; Vascular disease | ALS | Yes |
| 13 | 73 | White | M | FTLD-TDP-43 inclusions with MND; Vascular disease. | FTD | Yes |
| 14 | 78 | White | M | TDP-43 and ubiquitin-positive inclusions in frontal, temporal, caudate and spinal cord; Degeneration of lateral and ventral cortico-spinal tract; Alzheimer's Braak III. | FTD | Yes |
| 15 | 43 | White | M | TDP-43-positive neurites; Neuronal loss; Purkinje loss; Spinal degeneration | FTD | Yes |
| 16 | 86 | White | M | Degeneration of lateral cortico-spinal tracts and ventral roots; loss of anterior horn cells in spinal cord; TDP-43-positive neurites in hippocampus, midbrain and frontal cortex. | FTD | Yes |
| 17 | 79 | Asian | M | MND; FTLD-TDP-43; atypical CTE with prominent subcortical and subcerebellar white matter degeneration; Alzheimer's Braak II/VI | FTD & Respiratory failure | Yes |
| 18 | 53 | Hisp/Latino | M | FTLD-TDP-43 with MND; Spinal degeneration. | FTD & Respiratory failure | Yes |

| Cohort 2: Guamanian Samples | | | | | | |
|---|---|---|---|---|---|---|
| 1 | 57 | Guam | F | No NFT | Non-neurological | No |
| 2 | 43 | Guam | F | No NFT | Non-neurological | No |
| 3 | 48 | Guam | M | Anoxic encephalopathy; Cardiac arrhythmia. | Focal acute myocardial infarction. | No |
| 4 | 49 | Guam | M | Probable corona insufficiency | Non-neurological | No |
| 5 | 45 | Guam | M | Muscle atrophy; Bulbar palsy; Dysarthria; Dysphagia; No dementia. | Guamanian-ALS | Yes |
| 6 | 34 | Guam | F | Muscle atrophy; Bulbar palsy | Guamanian-ALS | Yes |
| 7 | 54 | Guam | F | Marked atrophy of facial, messeter and temporal muscles; Respiratory dysfunction; Fixed palate and increased Jaw Jerk. | Guamanian-ALS | Yes |
| 8 | 47 | Guam | F | Bular atrophy; Spinal spasticity; Muscle atrophy | Guamanian-ALS | Yes |

**Supplementary Table S2:** List of oligonucleotides used in this study.

| **Primers** | **Gene** | **Nucleotide sequence 5′ - 3′** | **Purpose** |
|---|---|---|---|
| TV140-F | POLB | AGTGGGCTGGATGTAACCTG | SA-PCR |
| TV141-R | POLB | CCAGTAGATGTGCTGCCAGA | SA-PCR |
| TV161-F | POLB | CATGTCACCACTGGACTCTGC AC | LA-qPCR |
| TV162-R | POLB | CCTGGAGTAGGAACAAAAATT GCT | LA-qPCR |
| TV21 RPmix | NeuroD (Human) | RealTimePrimers.com | SA-PCR and qRT-PCR |
| H-NeuroD1-LA-F | NeuroD (human) | CCGCGCTTAGCATCACTAAC | LA-qPCR |
| H-NeuroD1-LA-R | NeuroD (human) | TGGCACTGGTTCTGTGGTATT | LA-qPCR |
| H-ENO2-LA-F | Enolase (human) | ACGTGTGCTGCAAGCAATTT | LA-qPCR |
| H-ENO2-LA-R | Enolase (human) | CCTGAAACTCCCCTGACACC | LA-qPCR |
| H-MyH2-LA-F | MyH2 (human) | AAAGCCTGCCAAGCCCTAAA | LA-qPCR |
| H-MyH2-LA-R | MyH2 (human) | TGGTCAGCATGGCAAGTGAA | LA-qPCR |
| H-MyH4-LA-F | MyH4 (human) | CAGGAGTGGTCCCTAAAGGC | LA-qPCR |
| H-MyH4-LA-R | MyH4 (human) | GTAAAAACACGGTCCCTGCC | LA-qPCR |
| H-Enolase Primer mix | Enolase (human) | RealTimePrimers.com | SA-PCR |
| H-MyH2 Primer mix | MyH2 (human) | RealTimePrimers.com | SA-PCR |
| H-MyH4 Primer mix | MyH4 (human) | RealTimePrimers.com | SA-PCR |

| | | | |
|---|---|---|---|
| POLR2A LA FP | POLR2A (RNAP II) | CCTGTCCACTAGCTACCCCT | LA-qPCR (11.3 kb) |
| POLR2A LA RP | POLR2A (RNAP II) | CCCCTCCCCCTAACAGATCA | LA-qPCR (11.3 kb) |
| POLR2A SA FP | POLR2A (RNAP II) | GCTGGGATCGTGAACGGTAG | SA-PCR/DNA-qPCR (0.295kb) |
| POLR2A SA RP | POLR2A (RNAP II) | ATGGCGACAACTCACCTGG | SA-PCR/DNA-qPCR (0.295kb) |
| DmCrebB LA F | CrebB Drosophila | TCGAGTGTCAGGTGTTACGG | LA-qPCR |
| DmCrebB LA R | CrebB Drosophila | TCGCCACTGTAGTGAAGAGG | LA-qPCR |
| DmCrebB SA F | CrebB Drosophila | AGATCCTGCGGAGTCTACGA | SA-PCR |
| DmCrebB SA R | CrebB Drosophila | GGCGAGGCTCACTAATCTGG | SA-PCR |
| DmNeurexin LA F | Neurexin Drosophila | GGTACACCGGACTGAATGGG | LA-qPCR |
| DmNeurexin LA R | Neurexin Drosophila | CTGGCTACACTTGGTGGGTC | LA-qPCR |
| DmNeurexin SA F | Neurexin Drosophila | TGCAACAACAGTGCCCTACT | SA-PCR |
| DmNeurexin SA R | Neurexin Drosophila | TAGCCTTAACGAGCGACCAC | SA-PCR |
